# CB2 activation during epileptogenesis recruits immunomodulatory monocytes and mitigates cognitive, affective and seizure outcomes

**DOI:** 10.64898/2026.08.30.748087

**Authors:** Wanda Grabon, Nadia Gasmi, Anne Ruiz, Anatole Lang, Béatrice Georges, Ophélie Hurtado, Victor Blot, Jacques Bodennec, Sylvain Rheims, Amor Belmeguenai, Laurent Bezin

**Author notes:** co-last (Amor Belmeguenai, Laurent Bezin).

## Abstract

Temporal lobe epilepsy (TLE) is frequently associated with severe cognitive impairment and psychiatric comorbidities that are highly disabling and not targeted by anti-seizure medications, highlighting the need for therapies that target epileptogenesis rather than merely suppress seizures. Despite strong evidence implicating neuroinflammation in this process, anti-inflammatory approaches have not yielded such therapies yet. Infiltrating peripheral monocytes are widely viewed as detrimental amplifiers of post–status epilepticus (SE) inflammation, although emerging data suggest context-dependent protective roles. The cannabinoid receptor type 2 (CB2), highly expressed in myeloid cells, represents a potential immunomodulatory target linking leukocyte recruitment and inflammatory polarization. Using a juvenile rat model of TLE, we characterized hippocampal neuroinflammation and CB2 expression during epileptogenesis and evaluated the effects of transient CB2 activation with the selective agonist GP1a. SE induced a rapid but transient inflammatory response and robust recruitment of peripheral monocytes that persisted as anti-inflammatory monocyte-derived macrophages. CB2 activation did not suppress early cytokine induction but selectively enhanced recruitment of anti-inflammatory monocytes and modestly increased anti-inflammatory signaling. Importantly, transient CB2 stimulation preserved synaptic plasticity, improved cognitive performance, reduced anxiety-like behavior, and delayed seizure onset. These findings identify CB2-dependent modulation of peripheral myeloid cells as a potential disease-modifying axis in TLE.

## 1. INTRODUCTION

Temporal lobe epilepsy (TLE) is the most frequent focal epilepsy, accounting for most drug-resistant cases and tightly associated with cognitive dysfunction, psychiatric comorbidities and major socio-economic burden (Pitkänen et al., 2015). Despite compelling evidence that neuroinflammation is a core driver of epileptogenesis in both experimental models and patients, there is still no therapy that prevents the transition from an epileptogenic insult to chronic epilepsy and its long-term sequelae (Aronica et al., 2017). Clinical imaging with translocator protein (TSPO) PET and analyses of surgical specimens demonstrate widespread, persistent glial activation in TLE, yet anti-inflammatory strategies have not translated into robust disease-modifying treatments, highlighting a gap between descriptive neuroinflammation and actionable therapeutic pathways (Aronica et al., 2017; Brackhan et al., 2016).

Experimental work has placed myeloid cells at the center of this gap. Following status epilepticus (SE), blood–brain barrier disruption and chemokine induction recruit circulating monocytes into the hippocampus, and preventing their brain entry attenuates neuroinflammation and limits early neurodegeneration and cognitive deficits (Tian et al., 2017; Varvel et al., 2016). These findings have shaped a prevailing paradigm in which infiltrating monocytes are regarded as uniformly detrimental amplifiers of post-SE inflammation, and in which preventing their recruitment is considered a promising antiepileptogenic strategy. However, peripheral immune profiling in TLE patients reveals complex leukocyte and cytokine signatures that are not easily reconciled with a purely pathogenic view of monocyte function, and are compatible with the existence of context-dependent protective programs (Vieira et al., 2016).

In other CNS pathologies, monocyte-derived macrophages (mo-mΦs) can indeed support repair. After stroke or spinal cord injury, infiltrating myeloid cells expressing Arg1 and other pro-resolving markers contribute to debris clearance, promote resolution of inflammation and are required for long-term functional recovery, whereas blocking their recruitment abolishes these benefits (Greenhalgh et al., 2016; Wattananit et al., 2016). Conceptual and experimental work now distinguishes resident microglia from mo-mΦs as non-redundant populations with distinct temporal roles in injury and repair, challenging the idea that all brain-infiltrating monocytes are harmful (London et al., 2013; Silvin et al., 2023). The cannabinoid receptor type 2 (CB2) offers a mechanistic handle on this axis: CB2 is highly expressed by circulating myeloid cells and upregulated in reactive microglia, and in multiple neuroinflammatory models CB2 agonists reduce leukocyte–endothelium interactions, limit immune cell infiltration into the CNS and protect the blood–brain barrier (Grabon et al., 2023; Murikinati et al., 2010; Ramirez et al., 2012; Rom et al., 2013; Yoon and Grimsey, 2025). Together with data showing that CB2 signaling can shift microglia and macrophages towards anti-inflammatory, tissue-protective states (Braun et al., 2018; Grabon et al., 2023; Komorowska-Müller and Schmöle, 2020; Valeriano et al., 2025), this has consolidated the view of CB2 as a brake on leukocyte recruitment and an endogenous pro-resolving pathway. In the context of SE, the intuitive prediction is therefore that CB2 activation should blunt the acute inflammatory surge by restraining monocyte entry into the brain.

Our recent work in an adult pilocarpine SE rat model of TLE challenges part of this view by dissecting the endogenous roles of myeloid subsets. We showed that peripheral monocytes infiltrate the hippocampus in large numbers 24h after SE and persist for weeks as mo-mΦs, but on entry transiently adopt an anti-inflammatory, neuroprotective phenotype characterized by high IL-10, Arg1 and CD206 expression, whereas resident microglia are the main drivers of the early pro-inflammatory peak (Grabon et al., 2025). We also mapped CB2 expression in the mouse brain and found that CB2 is predominantly expressed by microglia under basal conditions, and is dynamically upregulated in microglia during neuroinflammation (Grabon et al., 2024, 2023). Together, these findings position CB2 at the intersection of monocyte recruitment and myeloid inflammatory state, and raise the possibility that manipulating CB2 during epileptogenesis could influence how peripheral myeloid cells engage with the inflamed brain in a context where infiltrating monocytes can be transiently protective.

Here we test this concept in a Li-pilocarpine SE model in juvenile rats that recapitulates spontaneous seizures and cognitive and anxiety-like comorbidities of TLE (André et al., 2003). We first define the temporal and cellular architecture of the neuroinflammatory response and CB2 expression during epileptogenesis, focusing on the respective contributions of microglia and infiltrating monocytes to pro- and anti-inflammatory signaling. We then selectively activate CB2 with the high-affinity, CB2-selective agonist GP1a restricted to the epileptogenic period and assess its impact on the neuroinflammatory status at the transcript level, monocyte recruitment, hippocampal synaptic plasticity, cognitive and anxiety-like behaviors, and the emergence of spontaneous seizures. Here we show that transient CB2 activation during epileptogenesis does not blunt the acute inflammatory surge but instead selectively enhances monocyte infiltration while improving long-term cognitive and epileptic outcomes, identifying CB2-dependent modulation of peripheral myeloid cell recruitment as an unexpected, tractable immunomodulatory axis in TLE.

## 2. METHODS

The full details of the methods are given in the supplementary material.

### Experimental design in animals

The experimental design is illustrated in Figure S1. Six distinct groups of animals were similarly subjected to Li-Pilo-SE at P21. Group 1 was used for characterization of the brain inflammatory response following SE and did not receive any treatment. Rats from groups 2 to 6 were treated i.p. with the CB2-specific agonist GP1a (Tocris) following SE according to the same protocol: 3mg/kg, diluted in vehicle (5% EtOH, 5%DMSO), at SE+3h, +1D, +2D, +3D, +4D, +7D, +10D, +14D. Eight rats from group 3 were treated i.p. according to the same schedule with another CB2 agonist, JWH-133 (Tocris) at a dose of 1.5 mg/kg diluted in the same vehicle. The doses of GP1a and JWH-133 were chosen according to the literature (Braun et al., 2018; Li et al., 2018; Tang et al., 2016; Zarruk et al., 2012).

### Group 1 – Inflammation characterization

30 rats were used to evaluate inflammatory profiles at transcript level 7h, 1 day (1D) and 9 days (9D) following SE in the hippocampus. Analyses were performed in rats subjected to SE (SE+7h, n=7; SE+1D, n=8, SE+9D, n=10) and in control rats (CTRL, n=5). 15 rats were used to characterize cellular inflammatory events at the histological level. Brains were collected 1D or 9D following SE after transcardiac perfusion with NaCl and paraformaldehyde (PFA) tissue fixation. 5 rats were used to evaluate inflammatory status of sorted microglia and infiltrating monocytes using flow cytometry. Brains were collected 1D following SE for subsequent tissue dissociation and cell sorting. Once sorted, the inflammatory profile of cell populations was evaluated by RT-qPCR.

### Group 2 – Effect of GP1a on molecular inflammation (RT-qPCR)

49 rats were used to evaluate inflammatory profiles at transcript level 7h, 1D and 9D following SE in rat hippocampus. Analyses were performed in rats subjected to SE treated with vehicle only (SE+Veh, n=6/time point), in rats subjected to SE treated with GP1a (SE+GP1a: n=6-7/time point) and in control rats (CTRL+Veh: n=6/time point).

### Group 3 – Effect of GP1a on cellular inflammation (Immunohistology)

29 rats were used to evaluate cellular inflammation at the histological level. Brains were collected 1D or 9D following SE. Analyses were performed on brain slices from rats subjected to SE treated with vehicle only (SE+Veh, n=6/time point), in rats subjected to SE treated with GP1a (SE+GP1a: n=6/time point) or with JWH-133 (SE+JWH-133: n=8) and in control rats (CTRL+Veh: n=5).

### Group 4 – Effect of GP1a on inflammatory profiles of microglia and monocyte-macrophages (Flow cytometry)

8 rats were used to evaluate inflammatory status of sorted microglia and monocyte-macrophages using flow cytometry. Brains were collected 1D following SE for subsequent tissue dissociation and cell sorting (CTRL+Veh, n=2; SE+Veh, n=3; SE+GP1a, n=3). Once sorted, the inflammatory profile of cell populations was evaluated by RT-qPCR.

### Group 5 – Effect of GP1a on behavior

A total of 40 rats were subjected to behavioral tests to determine the effect of GP1a treatment on anxiety levels, learning and spatial and recognition memory (12 control rats administered with the vehicle, "CTRL+Veh", 14 rats subjected to SE and administered with the vehicle, "SE+Veh", 14 rats subjected to SE that received GP1a treatment, "SE+GP1a"). Rats were subjected to the elevated zero-maze and water exploration tests (WET) to measure anxiety levels during the 3^rd^ week post-SE, the Morris Water Maze (MWM) test to measure spatial learning during the 4^th^ week post-SE followed by a probe test to assess spatial memory retention, and the reversal MWM (rMWM) test during the 5^th^ week post-SE to measure spatial learning flexibility. Finally, the animals were subjected to the Novel Object Recognition (NOR) test during the 8^th^ week post-SE. Handling-induced seizures were manually counted during the 2 weeks of MWM corresponding to the end of epileptogenesis.

### Group 6 – Effect of GP1a on hippocampal long-term potentiation

16 rats were used in electrophysiological studies to measure the effect of GP1a treatment on long-term potentiation in fresh hippocampal slices, monitored 3-4 weeks following SE (CTRL: n=6; SE: n=5; SE+GP1a: n=5).

### Group 7 – Effect of electrode implantation on monocyte infiltration

5 healthy rats were used to evaluate cellular inflammation at the histological level 1 month following surgical implantation of subdural screw electrodes and intracerebral electrode.

### Animals

Male Sprague Dawley rats were used in this study (Envigo, France). They were housed under a constant light/dark cycle (12/12h, lights on at 07:00 a.m.), with a room temperature kept at 23 ± 1°C. Animals arrived at 14 days of age (P14) and bred by groups of 10 with a foster mother until weaning. Following weaning, rats were maintained by groups of 2 with *ad libitum* access to food and water in standard cages. The experimental procedures were conducted in compliance with the European Community guidelines for care in animal research and approved by the CELYNE local Ethics Research Committee (protocol APAFIS#12194-2017041813592119 v2). Every effort was made to minimize animal suffering.

### Lithium-pilocarpine Status Epilepticus

SE was induced by pilocarpine, injected at P21. To prevent peripheral cholinergic side effects, scopolamine methylnitrate (1 mg/kg, s.c.; Sigma-Aldrich) was administered 30 min before pilocarpine hydrochloride (25 mg/kg, i.p.; Sigma-Aldrich). Lithium chloride (127 mg/kg, i.p.; Sigma-Aldrich) was injected 18 hours before scopolamine. After 30 min of continuous behavioral SE, 10 mg/kg diazepam was injected i.p., followed 90 min later, by a second injection of 5 mg/kg diazepam to terminate behavioral seizures. The animals were then treated with GP1a or vehicle as described in experimental design. At the end of the SE, the pups were returned to their foster mother and monitored until they resumed normal growth; they were therefore weighed daily.

### Brain collections

All rats were deeply anesthetized with isoflurane (4%) and administered with a lethal dose of pentobarbital (100 mg/kg; Euthasol). For RT-qPCR analysis (Groups 1 and 2), animals were intracardially perfused with ice-cold saline (30mL/min) to wash out blood from brain vessels. Hippocampi were rapidly microdissected on ice, frozen in liquid nitrogen, and stored at -80°C. For immunohistological analysis (Groups 1 and 3), animals were intracardially perfused with ice-cold saline and with 4% paraformaldehyde (PFA). For flow cytometry analysis (Group 4), animals were perfused intracardially with ice-cold saline (DPBS) supplemented with Actinomycin D (3µM) Anisomycin (100 µM) and Triptolide (10 µM) i.e. transcription and translation inhibitors. Hippocampi and the ventral limbic region (VLR), including the insular and piriform cortices, as well as the amygdala, were quickly dissected and placed in the same buffer until all perfusions were completed.

### Molecular biology

#### RNA extraction

Tissues (Groups 1 and 2) were crushed using Tissue-Lyser II (Qiagen) according to the manufacturer’s instructions. Total RNAs from brain structures were extracted using Tri-Reagent LS (Molecular Reasearch Center, #TS120). Genomic DNA was removed using Turbo-DNA-free (Ambion, #1907) and total RNAs were purified with RNeasy mini kit (Qiagen, #74104). For sorted cells (Group 4), total RNAs were extracted and purified using the RNeasy Plus Micro Kit (Qiagen, #74034) according to manufacturer’s instruction. RNA concentration was determined for each sample on the BioDrop® μLite.

### Reverse transcription and real-time quantitative PCR

Total tissue RNAs were reverse transcribed with PrimeScript RT Reagent Kit (Takara, #RR037A) according to manufacturer’s instructions. in the presence of synthetic external nonhomologous poly(A) standard messenger RNA (SmRNA) to normalize the RT step, as previously described (Grabon et al., 2024). Each cDNA of interest was amplified using the Rotor-Gene Q thermocycler (Qiagen), the SYBR Green PCR kit (Qiagen, #208052) and oligonucleotide primers (Eurogentec) specific to the targeted cDNA (**Table S1**). cDNA copy number detected was determined using a calibration curve, and results were expressed as cDNA copy number/µg tot RNA.

The pro-inflammatory index (PI-I) and anti-inflammatory index (AI-I) were calculated via a specific set of pro-inflammatory and anti-inflammatory genes, namely, IL-1β, IL-6, and TNFα, and IL-4, IL-10, and IL-13, respectively, via the formula given in the supplementary material (Grabon et al., 2024).

### Immunohistology

#### Tissue processing

Forty-micron-thick coronal sections were cut from frozen PFA-fixed brains via a cryomicrotome.

#### Immunolabelling

Detection of microglia was performed with goat anti-ionized calcium binding adaptor molecule 1 (Iba1) antibody (1:500, ab5076, Abcam). Infiltrating monocyte-macrophages (mo-MΦ) were labelled with mouse anti-Cluster of differentiation 68 (CD68) antibody (1:1000, MCA341GA, Bio-rad). Finally, astrocytes were double-detected with both mouse and rabbit anti-GFAP (1:1000, G3893, Millipore and 1:1000, ab5804, abcam respectively. The fluorescent secondary antibodies used are listed in the supplementary material. Nuclei were stained with DAPI (300 nM, Molecular Probes).

### Microscopy and analysis

Whole slices were scanned with a Carl Zeiss Axio Scan.Z1 Digital Slide Scanner (ZEISS) with a X20 lens on a 6 µm stack, using the pilot Zen (ZEISS). Images were then processed on with Fiji software (ImageJ).

### Flow cytometry

Samples were processed for tissue dissociation and cell sorting immediately following brain collection as quickly as possible. To prevent any artifactual *ex vivo* gene expression changes during brain dissociation and cell sorting procedures, all buffers and solutions used during the process (from animal perfusion to sorted cell flash freezing) were supplemented with the same cocktail of transcription and translation inhibitors as for brain collection (Ocañas et al., 2022).

### Brain tissue dissociation

Once collected, tissues were cut in smaller pieces with a scalpel and processed for dissociation using Miltenyi’s Adult Brain Dissociation Kit (#130-107-677) according to manufacturer’s instruction, running program 37C_ABDK_01. Inhibitor cocktail was added in each reagent.

### CD11b-positive cells magnetic enrichment

To increase fluorescence-activated cell sorting (FACS) yields and efficiency, cell suspensions were first enriched via the magnetic-activated cell sorting (MACS) technique, in which CD11b-positive cells (microglia and infiltrating monocytes) were magnetically separated for subsequent FACS via CD11b/c microbeads according to the manufacturer’s instructions (Miltenyi #130-105-634) from other cells before direct freezing. Details are provided in the supplementary material.

#### FACS

Microglia (CD11b^+^CD45^lo^CD11a^lo^) and infiltrating monocytes (CD11b^+^CD45^hi^CD11a^hi^) were sorted with a BD FACS Aria™ III Cell Sorter (BD Biosciences), as detailed in the supplementary material.

### Behavioral tests

All behavioral tests were carried out using equipment supplied by Noldus (Netherlands), which provided each test apparatus, camera system, infra-red lighting and tracking software (Ethovision XT video tracking system, Noldus, Netherlands). For all behavioral tests, the running order of animals was determined randomly. In the event of a seizure before or during the task, the rat was returned to its cage for at least 15 min before being retested.

### Elevated Zero-maze

The elevated zero-maze is a modification of the elevated plus-maze, suggested to have a higher sensitivity to measure anxiety level in rodent (Shepherd et al., 1994). The apparatus consists in an elevated ring 1m in diameter comprising two open portions and two portions protected by dark walls. The rats’ movements were recorded for 5 min by video tracking. Infra-red lamps and camera made it possible to measure the rats’ trajectories even in the dark areas protected by the walls.

### Water Exploration Test

As previously described, the WET further allows for the robust measurement of anxiety levels in rodent (Fares et al., 2013). A circular pool (180 cm in diameter) was filled with water maintained at 25°C to a depth of 40 cm and divided into four quadrants of identical size using opaque plastic panels to allow four rats to be recorded simultaneously. The rats’ trajectory was recorded for 5 min to measure total distance, average speed and time spent in the virtual central zone considered as anxiogenic.

### Morris Water Maze

Spatial learning in rats was tested over 5 days using the Morris Water Maze Test (Morris, 1984) in a 180 cm circular pool. The water was maintained at 25°C throughout the testing sessions. The tank was divided into 4 virtual quadrants: North (N), East (E), South (S) and West (W). A circular Plexiglas platform (14 cm in diameter) was hidden 1.5 cm below the water surface at a constant position in the north quadrant. The various spatial cues placed on the walls of the room around the pool were kept constant throughout the test. Details of the test are provided in supplementary material. A real-time video acquisition system was used to record platform-finding latency, distance and average swimming speed for each trial. The week following MWM spatial learning, retention was assessed during a "probe" session, after removing the platform. Rats’ trajectories were recorded for 90 sec to measure latency to platform ex-position and time spent in the target quadrant (N).

### Reversal Morris Water Maze

To measure spatial learning flexibility, the same experimental paradigm was used immediately afterwards, changing the position of the platform from the N to the S quadrant. On the first day of this second week, rats were positioned for 60 sec on the platform at its new location over two sessions, 3 hours apart. On the following 4 days, the rats were subjected to the same sessions as the MWM week, reversing the position of the starting points. The MWM reversal week was also followed by a 90-second "probe" session without a platform to measure latency to platform ex-position and time spent in both the target quadrant (S) and the previous target (N).

### Strategy analysis during MWM and Reversal MWM tasks

the Rtrack package was used for the analysis of tracking data, obtained during the MWM and the reversal MWM tasks (https://rupertoverall.net/Rtracl/index.html) with the recommended default parameters. Tracks have been classified in 8 different strategies, illustrated in **Figure S2**.

### Handling-induced seizures

The number of handling-induced seizures (Kouchi et al., 2022) was strictly reported during all the tests carried out with the Morris pool, i.e. a total of 30 sessions (15 sessions during the MWM test, 13 sessions during the MWM reversal and 2 probe sessions), as these tests were carried out at the end of the epileptogenesis period, i.e. 3-4 weeks following SE.

### Novel Object Recognition task

The NOR test is a widely used test for the investigation into memory alterations (Ennaceur and Delacour, 1988). Rats were first habituated to the empty apparatus, a 1 m^2^ arena with opaque sidewalls, during a 5 min session. The following day, during the familiarization session, the animals were presented with two identical objects (A and A’), positioned equidistant from the sidewalls. After a 4h delay, the animals were subjected to the testing session, presented with one familiar object (A) and one novel object (B) of similar size, placed in the same positions. To control for potential object-related bias, the identity of the objects (A and B) was counterbalanced across animals, such that each object served equally often as the familiar and the novel object. During both phases of the test, the exploratory behavior of rats was recorded with the videotracking system for a period of 5 min each. Animals were considered to be exploring an object when their nose was detected in the virtual 1 cm zone around the object.

### Electrophysiology

#### Slices preparation

Transverse hippocampal slices were prepared from P42-49 rats, i.e. 3-4 weeks following SE. Animals were anesthetized using isoflurane and sacrificed by decapitation. Brain was quickly extracted and cooled with ice-cold standard artificial cerebrospinal fluid (ACSF) bubbled with 95% O_2_ and 5% CO_2_. Hippocampi were dissected and 370 µm-thick transversal slices were prepared using a vibratome (VT1000S; Leica). The ACSF perfused during the recording was supplemented with 100 µM picrotoxin to block GABA_A_ receptors.

### Electrophysiological recordings

Whole-cell patch-clamp recordings were obtained from CA1 pyramidal neurons in current clamp mode at -70mV. Whole-cell patch clamp recordings were performed using an Axopatch-200B amplifier (Molecular Devices). Data were recorded and analyzed using a Digidata 1440A interface and pClamp 10 software (Molecular Devices). Stimulation at 0.05 Hz was used to establish baseline synaptic responses. The stimulation strength was set to evoke excitatory postsynaptic potentials (EPSPs) between 5 and 8 mV. Series resistance (typically 15-25 MΩ) was monitored throughout each experiment; cells with more that 20% change in series resistance were excluded from analysis.

### Long term potentiation

Once a steady baseline level of EPSPs was recorded for 10 to 20 min, long-term potentiation (LTP) was induced according to the Theta Burst Pairing (TBP) protocol at 5 Hz. A ten-trains sequence separated by 200 ms and composed of 5 pulses of 1 ms at 100 Hz coupled to backward action potentials (AP) is repeated 3 times at 10 seconds intervals. 0,05 Hz stimulations were resumed immediately after tetanization, and the amplitude of the EPSP recorded for 40 min. LTP is expressed as a percentage of increase in responses, relative to the baseline level previously recorded. Electrophysiological data were analyzed using pClamp 10 and Igor pro software (WaveMetrics).

### Stereotactic surgery

General anesthesia was induced by i.p. administration of a ketamine (80 mg/kg) and xylazine (10 mg/kg) mix, and local analgesia was induced by lidocaine (5 mg/kg; s.c.). Intracerebral electrode was positioned in the right basolateral amygdala. Subdural screw electrode was positioned over the right frontal cortex. The leads from both electrodes were inserted into a plastic pedestral and fixed using dental acrylic. Metacam (1 mg/kg, per os) was administered after surgery and for the following two days.

### Microscale thermophoresis

To check the affinity of GP1a for CB2, binding experiments were carried out by microscale thermophoresis (MST). Mouse mCB2-HIS-Tag was expressed and assembled into nanodiscs (Creative Biomart). The His-Tag domain was labelled using the Monolith Protein Labeling Kit RED-NHS according to the manufacturer’s instructions. mCB2-HIS-Tag (5 nM) was incubated 20 min with GP1a at 16 different concentrations obtained by serial dilution (from 62.5 nM to 1.9 pM) in buffer DPBS 0.01% pluronic. Samples were loaded into glass capillaries (Monolith NT Capillaries, Nanotemper Technologies), and thermophoresis analysis was performed using a Monolith NT.115 Series instrument (Nanotemper Technologies, excitation power 100%). Experiments were conducted as triplicates and treated with the MO.Affinity Analysis software (NanoTemper).

### Statistical Analysis

Statistical analyses were performed via Prism 10.0 software (GraphPad, USA), as detailed in the supplementary material. The results are presented as the mean ± SEM (standard error of the mean). Differences with a p-value<0.05 (p<0.05) were considered statistically significant. Details of the statistical tests for each figure are presented in the supplementary data (**Table S2**).

## 3. RESULTS

### 3.1 Characterization of the inflammatory response in the P21 rat Lithium-pilocarpine model

#### Inflammatory cytokine expression in the hippocampus

To characterize the early molecular inflammatory response after SE, we quantified prototypical cytokine transcripts in the hippocampus 7h, 1D and 9D post-SE (**Fig. 1**). IL-1β, IL-6 and TNFα transcripts showed a sharp, transient induction peaking at 7h (**Fig. 1A-C**, details of all statistical tests are provided in **Table S2**). IL-1β remained significantly different from controls at 1D and normalized by 9D, whereas IL-6 and TNFα were already back to control levels at 1D. The pro-inflammatory index (PI-I), computed from these three cytokines, similarly peaked at 7h and was no longer different from controls at 1D and 9D (**Fig. 1D**). In parallel, anti-inflammatory cytokines IL-4, IL-10 and IL-13 were transiently induced (**Fig. 1E-G**). IL-4 was significantly elevated at 1D, while IL-10 and IL-13 were increased at both 7h and 1D post-SE. The anti-inflammatory index (AI-I), based on these three cytokines, peaked at 7h and 1D and were at control levels at 9D (**Fig. 1H**).

**Figure 1.**
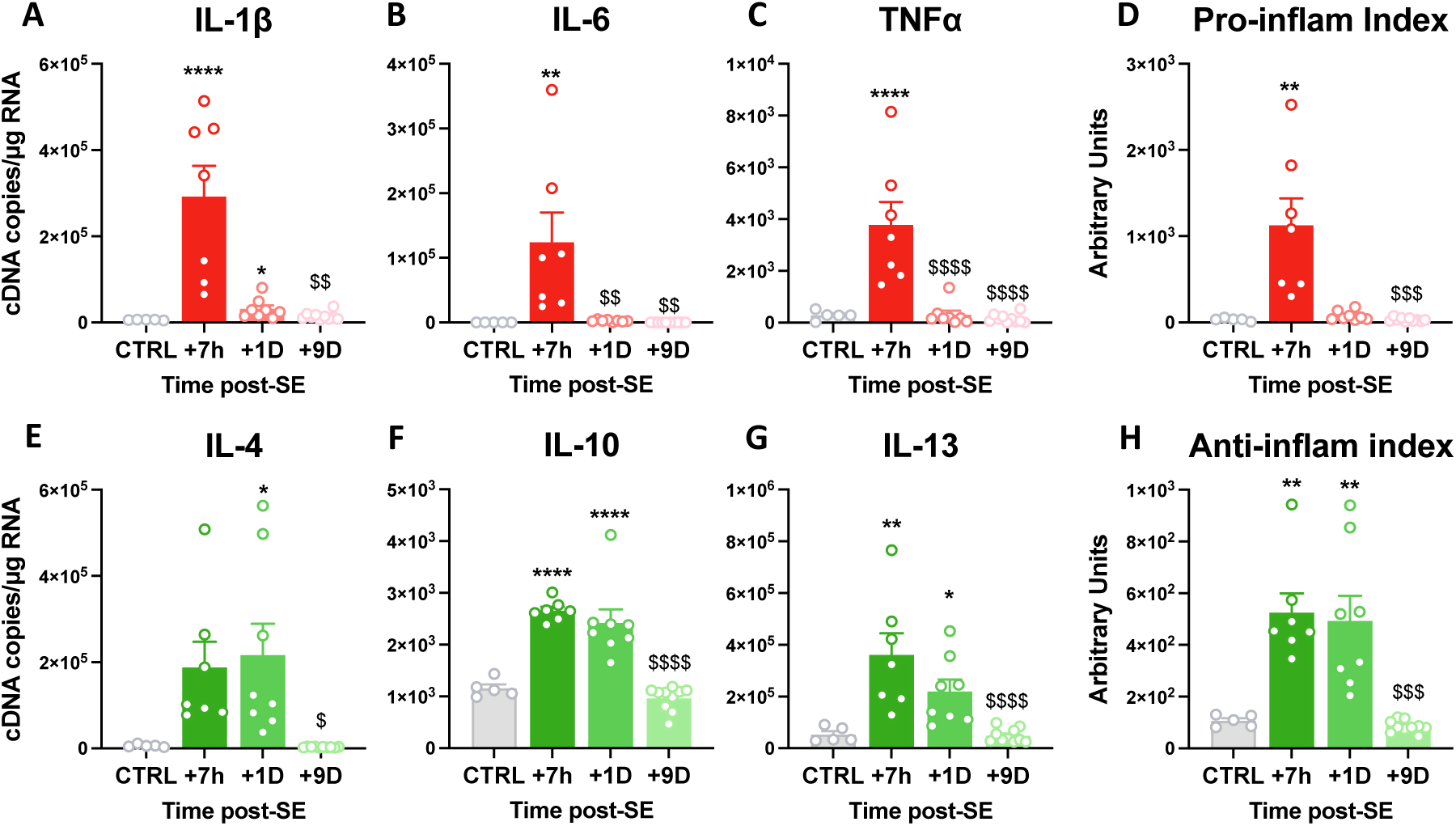
Early transient pro- and anti-inflammatory response in the hippocampus following SE induced by Li-Pilo at P21 in rats. Prototypical cytokine mRNA levels were quantified using calibrated RT-qPCR in hippocampi microdissected from brains of Group 1 rats (CTRL, n=5; SE+7h, n=7; SE+1D, n=8, SE+9D, n=10). Transcript levels are expressed as cDNA copy number per µg of total RNA. **A-C**. Quantification of transcript levels of pro-inflammatory cytokines IL-1β, IL-6 and TNFα transcript levels. **D**. Pro-inflammatory index was calculated as described in the Method section from IL-1β, IL-6 and TNFα and expressed in arbitrary units. **E-F**. Quantification of transcript levels of anti-inflammatory cytokines IL-4, IL-10 and IL-13. **H**. Anti-inflammatory index was calculated from IL-4, IL-10 and IL-13 transcript levels and expressed in arbitrary units. Normal data (IL-6, TNFα, IL-4, IL-10, IL-13 and AI-I) were analyzed with Tukey’s multiple comparison tests following one-way ANOVA, and non-normal data (IL-1β and PI-I) with Dunn’s multiple comparison tests following Kruskall-Wallis. All data are presented as mean + SEM. *: vs. CTRL; $: vs.SE+7h. */$, p<0.05; **/$$, p<0.01; ***/$$$, p<0.001; ****/$$$$, p<0.0001.

### Cellular inflammatory events triggered in the hippocampus following SE

To assess glial reactivity and monocyte recruitment after SE, we quantified transcript levels of astrocyte, microglial and monocyte markers and examined their cell morphology over time (**Fig. 2**). GFAP transcript levels were markedly increased at 1D and remained elevated at 9D post-SE, without detectable astrocytic morphological remodeling by immunolabelling (**Fig. 2A, E-G**). Microglia showed a transient threefold decrease in Iba1 transcripts at 7h, which normalized by 1D (**Fig. 2B**). Consistently, Iba1 protein almost disappeared from processes at 1D and became restricted to enlarged cell bodies at 9D (**Fig. 2K-M**). SE also induced robust monocyte recruitment, with MCP-1 mRNA levels upregulated more than 1500-fold at 7h and remaining high at 1D, preceding CD68 transcript induction (**Fig. 2C-D**). CD68-positive infiltrating monocytes with round morphology were detected in the hippocampus at 1D (**Fig. 2I-O**). By 9D post-SE, CD68 transcripts were maximal, monocyte-derived macrophages were more numerous, and some displayed a microglial-like morphology with CD68/Iba1 co-expression (**Fig. 2J, M, P**).

**Figure 2.**
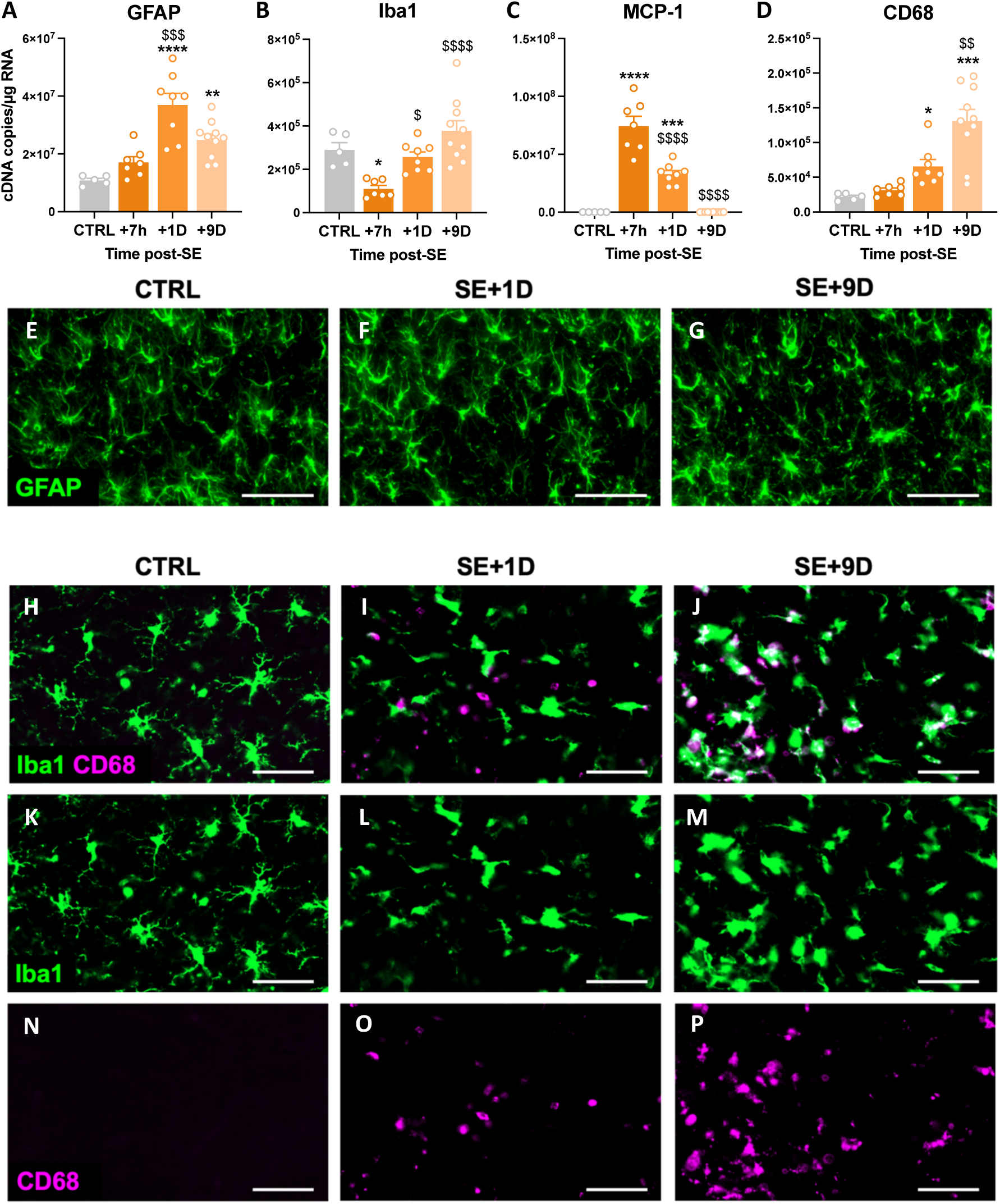
SE induced by Li-Pilo at P21 is followed by gliosis and monocyte infiltration in the hippocampus. **A-D.** Transcripts of cellular markers were quantified using calibrated RT-qPCR in hippocampi microdissected from brains of Group 1 rats (CTRL, n=5; SE+7h, n=7; SE+1D, n=8, SE+9D, n=10). Transcript levels are expressed as cDNA copy number per µg of total RNA. Quantification of GFAP (activation marker of astrocytes, **A**), Iba1 (microglial activation marker, **B**), MCP-1 (indicator of monocyte chemoattraction, **C**) and CD68 (monocytes and mo-mΦs marker, **D**) transcript levels. Details of statistical tests are presented in **Table S2**. All data are presented as mean + SEM. *: vs. CTRL; $: vs.SE+7h. */$, p<0.05; **/$$, p<0.01; ***/$$$, p<0.001; ****/$$$$, p<0.0001. **E-Q**. Immunodetection of astrocytes (GFAP, **E-G**), microglia (Iba1, **H-M**) and infiltrating monocytes (CD68, **H-J**, **N-P**) in the dentate gyrus of the hippocampus in control animals and in animals subjected to SE after 1D and 9D. Scale bars: 50µm.

### Comparative inflammatory profile of microglia and infiltrating monocytes 24h post-SE

To compare the inflammatory status of resident and infiltrating myeloid cells, we analyzed sorted microglia and monocytes/mo-mΦs 24h after SE (**Fig. 3A-D**). Monocytes/mo-mΦs (CD11b⁺CD45ʰⁱCD11aʰⁱ) and microglia (CD11b⁺CD45ˡᵒCD11aˡᵒ) were isolated by FACS after CD11b MACS enrichment from hippocampus and ventral limbic region (**Fig. 3A-B**). Tissue dissociation and sorting were performed in the continuous presence of transcription and translation inhibitors to limit *ex vivo* activation (Ocañas et al., 2022). At 24h, when the tissue pro-inflammatory peak is largely resolved but anti-inflammatory cytokines persist (**Fig. 1D, H**), monocytes/mo-mΦs did not show a higher inflammatory profile than microglia. TNFα transcripts were sixfold lower in monocytes/mo-mΦs, whereas IL-10 and typical M2 marker Arg-1 were 20- and 92-fold higher, respectively (**Fig. 3A-D**). Thus, infiltrating monocytes/mo-mΦs display a predominantly anti-inflammatory phenotype 24h after SE, consistent with observations in the adult P42 model (Grabon et al., 2025).

**Figure 3.**
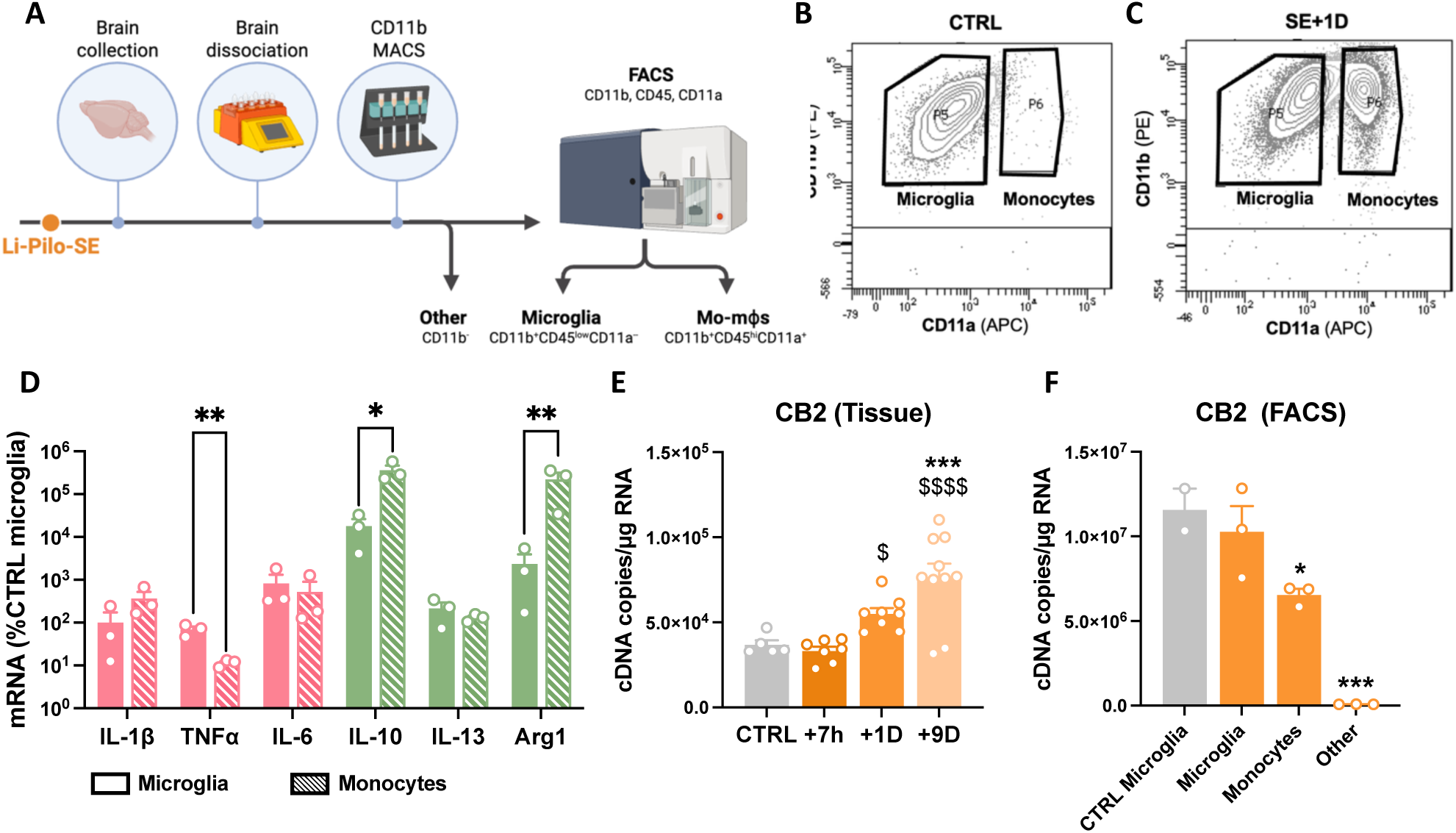
Infiltrating monocytes display a more anti-inflammatory phenotype than microglia and sustain increased CB2 expression in the hippocampus 24h post-SE. **A.** Workflow for microglia and monocyte-macrophages sorting. **B-C.** Microglia (CD11b^+^CD45^lo^CD11a^lo^) and infiltrated monocytes (CD11b^+^CD45^hi^CD11a^hi^) were sorted by FACS following CD11b MACS enrichment from hippocampus and VLR collected 1D post-SE in Group 5 rats (CTRL, n=2; SE+1D, n=3). **D**. Transcript levels of inflammatory markers were quantified by RT-qPCR in the sorted populations. and expressed as a percentage of the levels measured in the microglia of control rats. **E-F**. CB2 mRNA was quantified using calibrated RT-qPCR in hippocampi microdissected from brains of Group 1 rats (CTRL, n=5; SE+7h, n=7; SE+1D, n=8, SE+9D, n=10; **E**) and in cells sorted from hippocampus and VLR collected 24h post-SE in Group 5 rats (CTRL, n=2; SE+1D, n=3; **F**). Transcript levels are expressed as cDNA copy number per µg of total RNA. Details of statistical tests are presented in **Table S2**. Data are presented as mean + SEM. **E**: $: vs SE+7h; *: vs CTRL. **F**: *: vs. SE microglia. */$, p<0.05; **/$$, p<0.01; ***/$$$, p<0.001; ****/$$$$, p<0.0001.

### 3.2. CB2 as a target to modulate myeloid-driven neuroinflammation

CB2 is predominantly expressed by myeloid cells and has been implicated in the control of neuroinflammatory responses (Grabon et al., 2024, 2023), making it a relevant candidate to modulate the myeloid-driven inflammation identified in the P21 Li–Pilo model.

We then quantified its mRNA levels in hippocampal tissue and in sorted cell populations after SE (**Fig. 3A-C**). Tissue CB2 transcript levels increased in the hippocampus between 7h and 1D and reached a maximum at 9D post-SE (**Fig. 3E**). We focused our cellular analyses on 1D, when CB2 induction is already significant and monocyte infiltration is robust, allowing us to relate the tissue increase in CB2 to early infiltrating monocytes rather than to later established monocyte–macrophages. Infiltrating monocytes at 1D post-SE exhibit a less pro-inflammatory profile (lower TNFα) and a more anti-inflammatory profile (higher IL-10 and Arg1) than resident microglia (**Fig. 3D**). At this time point, CB2 was expressed in microglia and infiltrating monocytes, but not in CD11b-negative cells (**Fig. 3F**), in line with its predominantly restricted expression to myeloid cells in mouse brain (Grabon et al., 2024). At 1D, CB2 mRNA levels in microglia were similar in controls and rats subjected to SE, indicating that the tissue-level increase primarily reflects monocyte influx rather than induction in resident microglia.

To selectively activate CB2 in this context, we used the highly specific agonist GP1a and first validated its binding to CB2 by microscale thermophoresis (**Fig. S3**). Using mCB2-HIS-Tag assembled into nanodiscs, GP1a generated clean MST signals without autofluorescence, aggregation or photobleaching artefacts (**Fig. S3A-B**), and yielded a dissociation constant (K_d_) of 1.19 ± 0.52 nM (**Fig. S2C**), confirming strong binding to CB2.

In all subsequent experiments, GP1a was administered starting 3h after SE onset, i.e. before the inflammatory peak, and continued for 2 weeks (1, 2, 3, 4, 7, 10 and 14 days post-SE) to cover the epileptogenic period exclusively, at distance from epilepsy onset (4–7 weeks post-SE). Weight regain in the days following SE is a valuable indicator of recovery (Turski et al., 1989). Body weight was monitored in rats included in experiments lasting ≥ 6 days (parts of Groups 2, 3, 5 and 6), yielding 23 healthy controls given vehicle (CTRL+Veh), 26 SE rats given vehicle (SE+Veh) and 31 SE rats treated with GP1a (SE+GP1a) (**Fig. S4A–C**). Body weight at P21 (SE induction) was similar across groups (**Fig. S4A**). As expected, SE induced a marked weight loss at D1 (**Fig. S4C**). SE+GP1a rats lost slightly less weight than SE+Veh rats and showed a significantly greater weight gain by day 6 post-SE (**Fig. S4B–C**).

### 3.3. GP1a treatment during epileptogenesis reshapes hippocampal neuroinflammation

#### Early cytokine response to GP1a treatment

We first examined whether GP1a treatment altered the early hippocampal cytokine response to SE. CB2 transcript levels were increased in SE+Veh rats at 1D and 9D compared with CTRL+Veh, and a single GP1a administration was sufficient to raise CB2 mRNA already at 10 h, to nearly twice the level measured in SE+Veh rats at this time point (**Fig. 4A**). The pro-inflammatory index (PI-I), calculated from IL-1β, IL-6 and TNFα, was transiently elevated 10h after SE and was not significantly reduced by GP1a (**Fig. 4B,D–F**). In contrast, the anti-inflammatory index (AI-I), based on IL-4, IL-10 and IL-13, was significantly higher in SE+GP1a than in SE+Veh rats at 10h, largely driven by an over-induction of IL-4 transcripts, while no differences between the two groups persisted at 1D or 9D (**Fig. 4C,G–I**).

**Figure 4.**
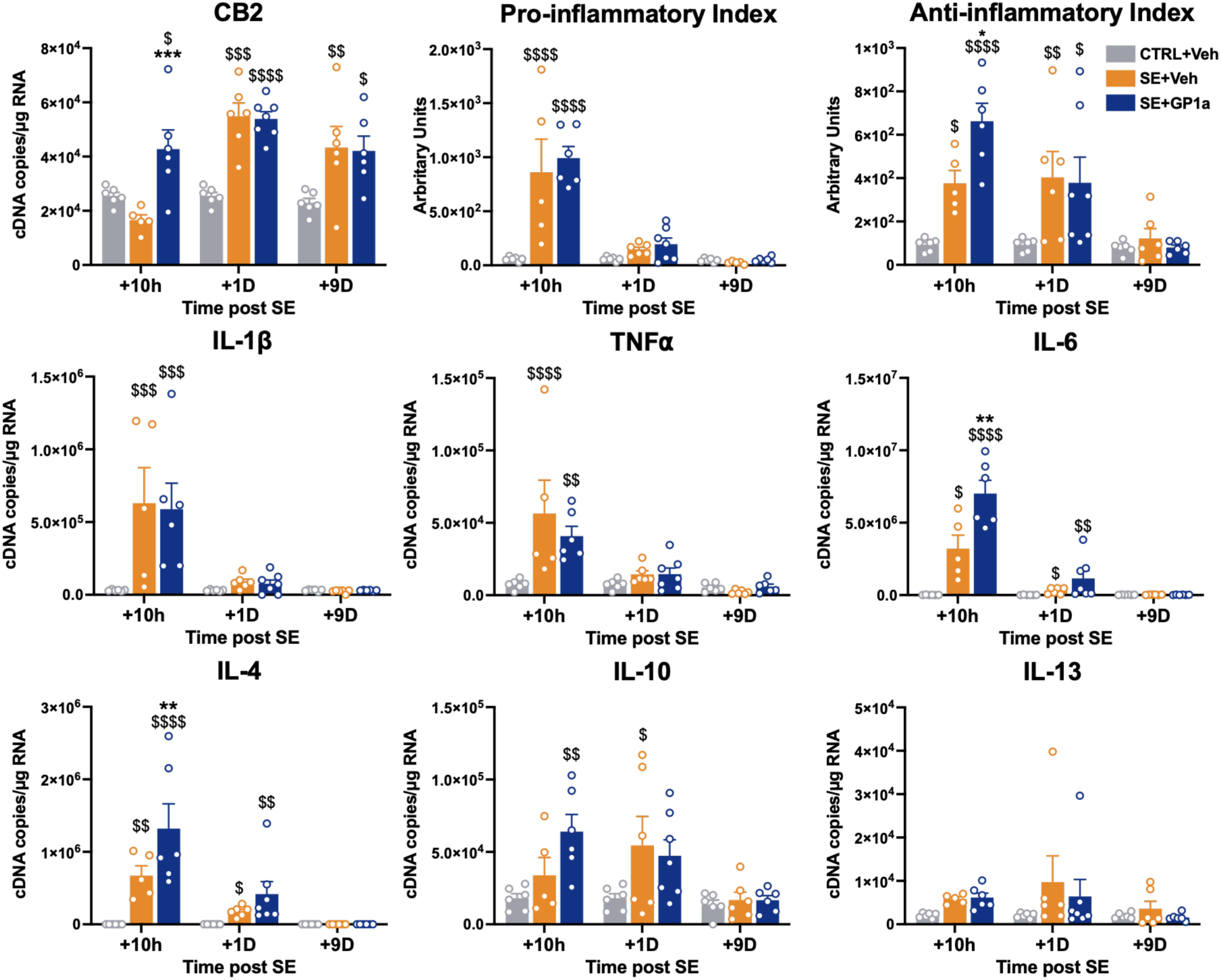
GP1a treatment midly modulates early inflammatory cytokines expression in the hippocampus. CB2 (**A**) and prototypical inflammatory cytokine (**D-I**) mRNA was quantified using calibrated RT-qPCR in hippocampi microdissected from brains of Group 2 rats (CTRL+Veh, n=6/time point; SE+Veh, n=5-6/time point; SE+GP1a; n=6-7/time point) collected 10h, 1D or 9D following SE. **A**. Quantification of CB2 transcript level. **B-C**. Pro-and anti-inflammatory index were calculated as described in the Method section from IL-1β, TNF*-*α and IL-6 and from IL-4, IL-10 and IL-13 respectively*. **D-I.*** Quantification of inflammatory cytokine transcript level are expressed as cDNA copies per µg of total RNA and presented as mean + SEM. Details of statistic tests are presented in Table S2. $: vs CTRL+Veh; *: SE+Veh vs SE+GP1a. */$, p<0.05; **/$$, p<0.01; ***/$$$, p<0.001; ****/$$$$, p<0.0001.

### GP1a does not modify astrocyte or microglia activation

The effect of CB2 activation on cellular inflammatory events following SE was then assessed by quantifying GFAP and Iba1 transcript levels and by immunohistochemistry at 1D and 9D post-SE. GP1a treatment did not alter GFAP mRNA levels or astrocyte morphology in the hippocampus at either time point, nor did it significantly affect Iba1 expression or microglial morphology (**Fig. S5A–K**).

### CB2 activation with GP1a or JWH-133 promotes monocyte infiltration

To assess the effect of GP1a on the infiltration of circulating monocytes, we first quantified MCP-1 expression in hippocampal tissue 10h, 1D and 9D after SE (**Fig. 5A**). In SE+GP1a rats, MCP-1 transcript levels were about 1.5-fold higher than in SE+Veh rats 10h after SE. Infiltrating monocytes were then visualized and quantified by CD68 immunodetection on coronal hippocampal sections collected 1D and 9D after SE (**Fig. 5B–E**). Very few CD68-positive cells were detected in healthy brains. At 9D post-SE, GP1a treatment markedly increased CD68-positive cell number and area in the hippocampus, to more than twice the values measured in SE+Veh rats, with a particularly strong effect in CA1 (**Fig. 5B–E**). Because CB2 activation is often reported to reduce monocyte infiltration in other neuroinflammatory models (Braun et al., 2018; Chung et al., 2016; Sheng et al., 2019), we repeated the experiment using the CB2 agonist JWH-133. A new cohort of pups underwent Li–pilocarpine SE and received either vehicle or JWH-133 (1.5 mg/kg, i.p.) according to the same schedule as GP1a. Nine days post-SE, JWH-133 also increased CD68-positive cell density and CD68-positive area in the hippocampus, to a degree comparable to that observed with GP1a (**Fig. 5G–H**).

**Figure 5.**
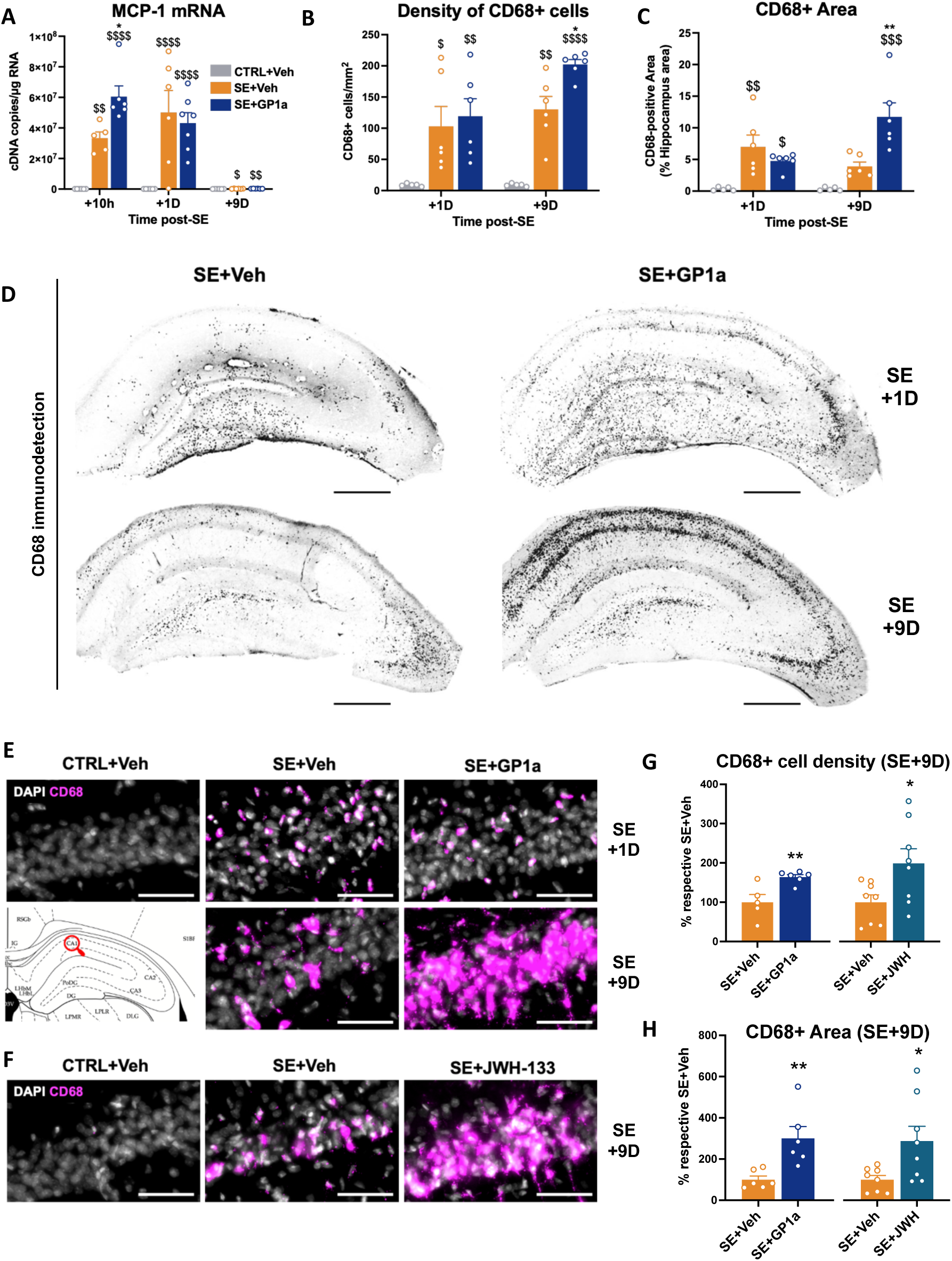
CB2 activation with either GP1a or JWH-133 treatment robustly promotes monocytes brain infiltration following SE. **A**. Transcript levels of MCP-1, involved in circulating monocyte chemotaxis to the injured brain, were quantified by RT-qPCR in the hippocampus 10h, 1D and 9D following SE collected from Vehicle-treated (SE+Veh, n=5-6/time point) and GP1a-treated SE rats (SE+GP1a, n=6/time point) and compared with healthy controls (CTRL+Veh, n=5), in Group 3 rats. **B-E**. Infiltrating monocytes were immunolabeled with anti-CD68 antibody (Biorad MCA341GA). CD68-positive surface area was quantified in dorsal hippocampus at bregma -3.30mm 1D and 9D following SE in Vehicle-treated (SE+Veh, n=6/time point) and GP1a-treated SE rats (SE+GP1a, n=6/time point) and compared with healthy controls (CTRL+Veh, n=5), in Group 4 rats. **B**. Quantification of CD68-positive cell density, measured in the whole hippocampus, expressed as CD68-positive cells/mm^2^. **C**. Quantification of CD68-positive surface area, measured in the whole hippocampus, expressed as percent of measured area **D**. Representative histological images of CD68 immunodetection, acquired with slide scanner in the hippocampus (x20 objective, scale bars: 500µm). **E**. Zoom of CD68 immunodetection in CA1 (scale bars: 50µm). **F-H**. Effect of CB2 activation on hippocampal mo-mΦs infiltration after SE was investigated following JWH-133 treatment (1.5 mg/kg, ip). Brains were collected 9D following SE (SE+Veh, n=8; SE+JWH-133, n=8) and mo-mΦs were labeled in the hippocampus with anti-CD68 antibody. **F**. Representative histological images of CD68 immunodetection following JWH-133 treatment in CA1, taken with slide scanner in the hippocampus (x20 objective, scale bars: 50 µm). CD68-positive cell density (**G**) and quantification of CD68-positive area (**H**) measured in the whole hippocampus 9D post-SE in both SE+GP1a and SE+JHW-133 rats expressed as % of respective controls (SE+Veh). Details of statistic tests are presented in supplementary data. Results are presented as mean + SEM. $: vs CTRL+Veh; *: SE+Veh vs SE+GP1a. */$, p<0.05; **/$$, p<0.01; ***/$$$, p<0.001; ****/$$$$, p<0.0001.

### Anti-inflammatory phenotype of infiltrating monocytes is preserved under GP1a treatment

To better characterize the role of infiltrating cells, monocytes (CD11b⁺CD45ʰⁱCD11aʰⁱ) and microglia (CD11b⁺CD45ˡᵒCD11aˡᵒ) were sorted by FACS after CD11b MACS enrichment from hippocampus and ventral limbic region 24h after SE, when monocyte numbers were high in both SE+Veh and SE+GP1a rats (**Fig. S6A–C**). CD11b⁺CD45ʰⁱCD11aʰⁱ cells from control rats, presumably perivascular macrophages, were too scarce for transcript analyses and were therefore excluded. GP1a treatment did not significantly affect the inflammatory status of any sorted population at 1D, but infiltrating monocytes consistently displayed an expression profile skewed towards an anti-inflammatory, M2-like phenotype compared with microglia (**Fig. S6D–I**)(Grabon et al., 2025).

### 3.4. GP1a treatment during epileptogenesis mitigates epilepsy-associated comorbidities and seizure occurrence

Temporal lobe epilepsy is frequently associated with cognitive deficits and affective comorbidities, including anxiety, which markedly affect quality of life and is worsened by currently available antiepileptic drugs (Khalid et al., 2024; Mazarati et al., 2017; Schmidt and Löscher, 2005). We therefore examined whether GP1a treatment during epileptogenesis mitigated seizure occurrence and epilepsy-associated comorbidities in the Li–pilocarpine P21 rat model.

*In vivo* and *ex vivo* tests were used to measure the effects of GP1a on cognitive dysfunctions at the end of epileptogenesis. Group 5 animals (CTRL+Veh, n=12; SE+Veh, n=14; SE+GP1a, n=14) were subjected to behavioral tasks to assess learning and memory skills *in vivo*, and hippocampal slices from Group 6 animals (CTRL, n=6; SE, n=5; SE+GP1a, n=5) were used to measure CA1 long-term potentiation (LTP), thereby evaluating the impact of GP1a treatment during epileptogenesis on hippocampal synaptic function.

### Spatial learning and memory

Spatial learning abilities were tested using the MWM during the 3rd week post-SE, cognitive flexibility was tested with the reversal MWM (rMWM) during the 4th week post-SE by changing the platform position, and retention was measured using probe tests after platform removal at the end of each week (**Fig. 6A–I**).

**Figure 6.**
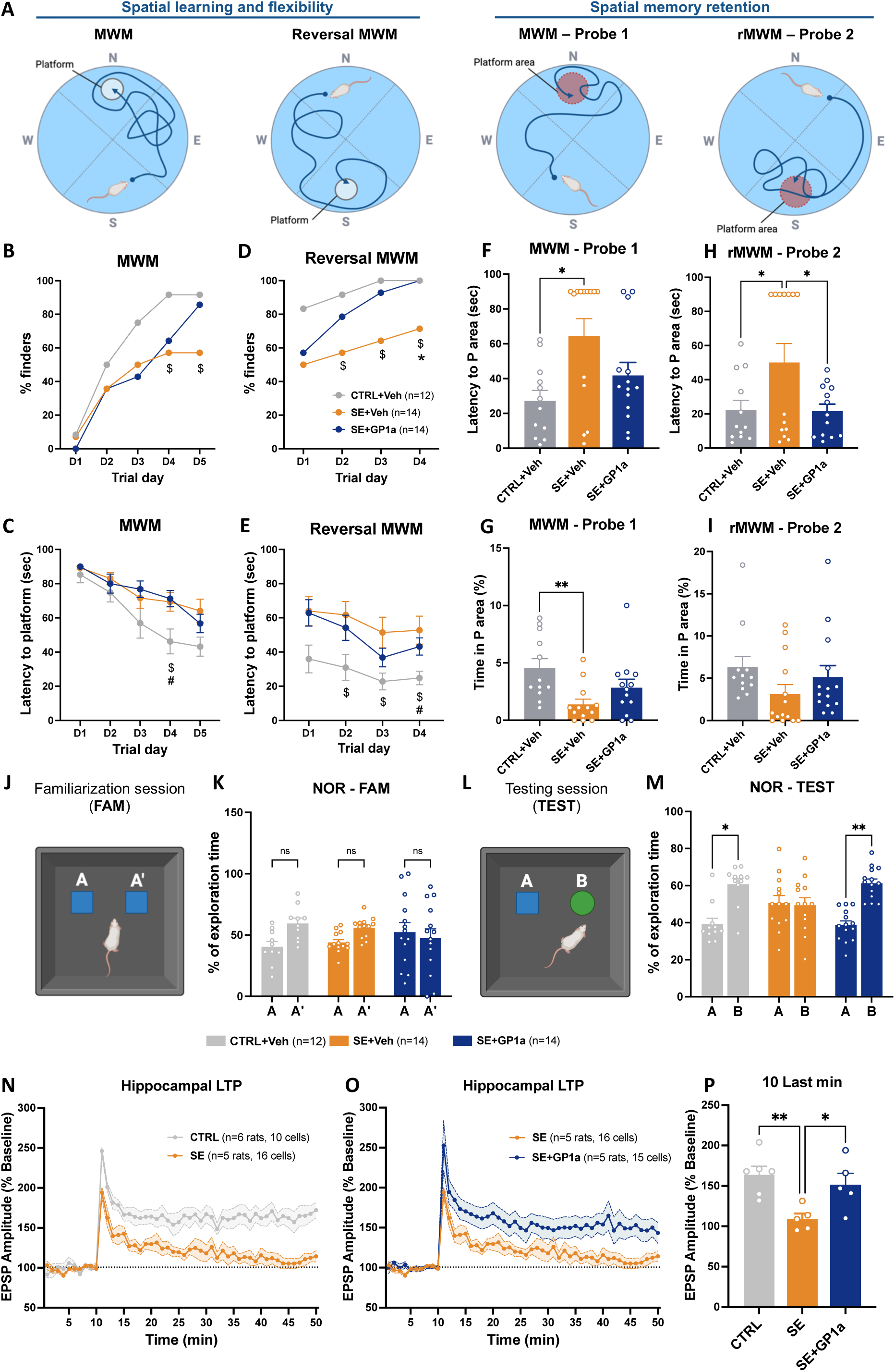
GP1a treatment during epileptogenesis protects cognitive functions. Group 5 animals (CTRL+Veh, n=12; SE+Veh, n=14; SE+GP1a, n=14) were subjected to behavioural tasks to assess GP1a effect on learning and memory skills *in vivo*. Hippocamps slices from Group 6 animals (CTRL, n=6; SE, n=5; SE+GP1a, n=5) were used to measure Long Term Potentiation (LTP) in CA1 pyramidal neurons to evaluate GP1a effect on synaptic plasticity at the cellular level. **A-I. Morris Water Maze**. **A**. Positions of the hidden platform or the platform area (P area) in each task. **B-C**. Rats were subjcted to the MWM during the 3^rd^ week post-SE. The number of rats that found the platform at least once during the day (% finders) was counted each trial day (**B**). Latency, i.e. time to reach the platform, was measured on each trial (sec), and averaged over each trial day ; maximum trial duration was 90 sec (**C**). **D-E**. The following week, rats were subjected to the rMWM task. % of finders (**D**) and latency to platform (**E**) were measured as for the MWM task. $: CTRL+Veh vs SE+Veh; #: CTRL+Veh vs SE+GP1a; *: SE+Veh vs SE+GP1a. **F-I**. Following MWM and rMWM, rats were subjected to probe tests after platform removal. Latency to P area was measured (**F, H**) together with time spent in this area (**G, I**). **Novel Object Recognition**. Rats were subjected to the NOR task during the 8^th^ week post-SE. **J**. Illustration of the familiarization session. **K**. Time spent exploring objects A and A’ was measured during the task and expressed as % total exploration time for each rat. **L**. Illustration of the testing session, 4h later. **M**. Time spent exploring familiar object A and novel object B was expressed as % total exploration time. **N-P. Hippocampal LTP.** Hippocampal LTP elicited by Theta-burst pairing (TBP) in CA1 pyramidal neurons was monitored during the 3-4^th^ week post-SE. 2-4 cells were analyzed for each rat. Excitatory Post Synaptic Potential (EPSP) was recorded for 10 min as a baseline before TBP tetanization and 50 min after to mesure LTP. At each time point, the values obtained for cells from the same rat were averaged, then for each group, the values obtained for each rat were averaged. LTP measured in SE animals was compared with that of healthy rats (**N**) and GP1a-treated SE rats (**O**). **P**. Mean EPSP amplitude was calculated over the last 10 min. Results are presented as mean of rats + SEM. Details of statistic tests are presented in supplementary data. Ns, non-significant; *, p<0.05; **, p<0.01.

#### Morris Water Maze

During the MWM task, CTRL+Veh rats successfully learned the hidden platform position, as the proportion of “finders” (rats that found the platform at least once during the day) increased from 8.3% on day 1 to 91.7% on day 5, and escape latency significantly decreased over time (**Fig. 6A–C**). In SE+Veh rats, only 57.1% had found the platform by the end of the week; their mean latency decreased over training but remained significantly higher than in CTRL+Veh on day 4 (**Fig. 6A–C**). SE+GP1a rats were more numerous to find the platform than SE+Veh rats by the end of the week, although this difference did not reach significance (p=0.0943), and their mean latency decreased across days but was not significantly lower than that of SE+Veh rats (**Fig. 6A–C**).

#### Reversal Morris Water Maze

On the first day of rMWM, rat performance in each group remained closed to that observed on the last day of MWM (**Fig. 6D–E**). By the end of the rMWM week, 100% of SE+GP1a rats had found the platform at least once during the day, compared with 71.4% of SE+Veh rats (p=0.0308, **Fig. 6D**). Latency to the hidden platform was significantly higher in SE+Veh rats than in CTRL+Veh rats on trial days 2, 3 and 4 and did not significantly improve over the four days, whereas latency in SE+GP1a rats significantly decreased over the same period (**Fig. 6E**).

#### Probe tests

Spatial memory was measured after each training week (Probe 1 after MWM and Probe 2 after rMWM) by removing the platform and quantifying both the latency to first visit to the former platform area (P area) and the cumulative time spent in P area. In Probe 1, SE+Veh rats were significantly slower than CTRL+Veh rats to reach P area and spent less time in P area (**Fig. 6F–G**). SE+GP1a rats showed intermediate values for both measures, which were not statistically different from either CTRL+Veh or SE+Veh rats (**Fig. 6F–G**). In Probe 2, SE+Veh rats were again slower than CTRL+Veh rats to reach P area, while SE+GP1a rats were significantly faster than SE+Veh rats to reach P area (**Fig. 6H**).

#### Search strategy in the water maze

Beyond conventional MWM performance measures, swim paths were analyzed and classified into non-goal oriented, procedural and allocentric strategies using Rtrack. All groups of rats showed a comparable learning-related transition from non-goal-oriented to more efficient spatially guided strategies across acquisition sessions, with no significant group differences. However, during Probe Trial 2 (Reversal MWM), SE+Veh rats exhibited increased thigmotaxis, a wall-oriented, non-goal-oriented swimming strategy commonly associated with anxiety-like behavior. This increase was significantly reduced by GP1a treatment, consistent with an attenuation of anxiety-related behavioral component associated with SE (**Fig. S2C**).

### Recognition memory

Recognition memory was tested using the Novel Object Recognition (NOR) task during the 8th week post-SE, i.e. during the chronic phase and at a late time point after the last GP1a or vehicle administration (**Fig. 6J–M**). During the familiarization session, the three groups spent a similar amount of time exploring the two identical objects A and Aʹ (**Fig. 6J–K**). Four hours later, rats were presented with a familiar object (A) and a novel object (B) in the same arena (**Fig. 6L**). CTRL+Veh rats correctly memorized the familiar object, as they spent more time exploring the novel object B, in line with intact recognition memory (Ennaceur and Delacour, 1988) (**Fig. 6M**). SE+Veh rats did not explore the novel object B more than object A, indicating impaired recognition memory, whereas SE+GP1a rats, like CTRL+Veh rats, spent more time exploring the novel object B than the familiar object A, indicating preserved recognition memory (**Fig. 6M**).

### Hippocampal long-term potentiation

Hippocampal LTP is a well-established candidate mechanism for learning and memory (Bliss and Collingridge, 1993). It was elicited in CA1 pyramidal neurons by a theta-burst pairing (TBP) protocol and measured during the 3rd–4th week post-SE in Group 6 rats, i.e. at the same time as Group 5 rats were tested in the MWM (**Fig. 6N–P**). LTP was significantly reduced in SE rats compared with CTRL rats (**Fig. 6N–O**). Mean excitatory postsynaptic potential (EPSP) amplitude over the last 10 minutes of recording was 164.8 ± 10.4% of baseline in CTRL rats and 109.3 ± 6.5% in SE rats (p=0.008), confirming impaired synaptic plasticity after SE (**Fig. 6P**). GP1a treatment during epileptogenesis protected LTP, as EPSP amplitude in SE+GP1a rats reached 150.5 ± 13.9% of baseline and was significantly higher than in SE rats (**Fig. 6P**).

### Anxiety-like behavior

Behavioral tests were used to measure the effects of GP1a on anxiety-like behavior at the end of epileptogenesis. Rats were subjected to the elevated zero-maze (O-maze) and the Water Exploration Test (WET) at the end of the second week post-SE (**Fig. 7**).

**Figure 7.**
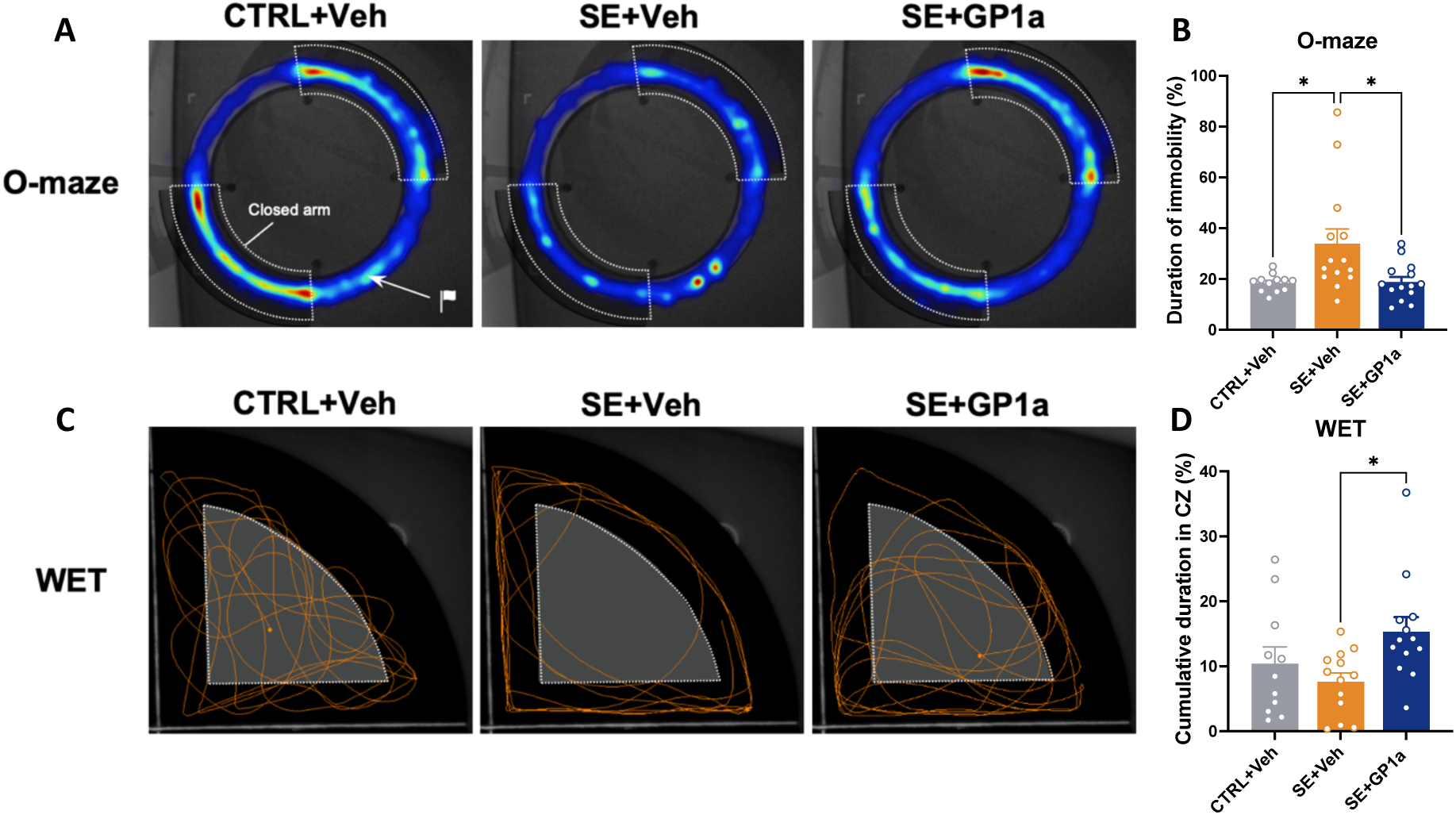
GP1a treatment during epileptogenesis reduced anxiety-like behavior. Group 5 animals (CTRL+Veh, n=12; SE+Veh, n=14; SE+GP1a, n=14) were subjected to Elevated zero-maze (O-maze) and Water Exploration Test (WET) at the end of the second week post-SE. **A-B. O-maze. A**. Overlay of heatmaps by group, generated by the videotracking software during the O-maze task (Ethovision X, Noldus). The regions explored are colored. The warmer the color, the more the region has been explored on average by the group; the colder it is, the less time the animals have spent there. **B**. Immobility duration was quantified over the entire test duration (5 min) and expressed as a % of total time. **C-D**. **WET**. **C**. Representative tracks of one rat of each group. **D**. Time spent in central zone (CZ) was quantified over the entire test duration (5 min) and expressed as a % of total time. O-maze and WET results were analyzed with Dunn’s multiple comparison test following Kruskall-Wallis’ test. Results are presented as mean + SEM, *, p<0.05.

#### O-maze

The elevated zero-maze is used to assess anxiety in rodents by quantifying time spent in open arms (OA), which are considered anxiogenic (Shepherd et al., 1994). As shown by representative heatmaps, CTRL+Veh rats spent considerable time near the exits of the closed arms and explored the open arms to some extent, whereas SE+Veh rats displayed less exploratory behavior and tended to freeze at the starting point (**Fig. 7A**). GP1a-treated SE rats showed an exploration pattern visually similar to CTRL+Veh rats (**Fig. 7A**). Because SE+Veh rats frequently froze in the OA, time spent in OA could not be reliably quantified; we therefore measured cumulative immobility time regardless of the zone (**Fig. 7B**). SE+Veh rats spent significantly more time immobile than CTRL+Veh rats, whereas SE+GP1a rats showed significantly lower immobility time than SE+Veh rats and values not different from CTRL+Veh rats (**Fig. 7B**).

#### Water Exploration Test

As previously described, the WET provides a robust measure of anxiety in epileptic rodents by quantifying time spent in the central zone (CZ) of a quadrant of the pool, considered anxiogenic (Fares et al., 2013). In this task, the more time a rat spends in the CZ, the less anxious-like its behavior is. Representative tracks illustrate reduced central exploration in SE+Veh rats compared with CTRL+Veh rats, whereas SE+GP1a rats showed greater use of the CZ (**Fig. 7C**). On average, SE+GP1a rats spent about twice as much time in the CZ as SE+Veh rats (**Fig. 7D**).

### Epileptogenesis

In a preliminary study, healthy rats were implanted with subdural screw and intracerebral electrodes to assess post-surgical inflammation, particularly monocyte infiltration. CD68 immunodetection one month after surgery revealed substantial numbers of CD68-positive cells, especially in neocortex near the implantation sites, and to a lesser extent in hippocampus and thalamus (**Fig. S7**). To avoid any inflammatory confound induced by electrode implantation, the effect of GP1a on seizure onset was therefore evaluated only at the behavioral level by counting handling-induced seizures, as previously reported (Kouchi et al., 2022).

Handling-induced seizures were counted manually, regardless of severity, based on observed behavioral manifestations during the two weeks of MWM testing, corresponding to the end of epileptogenesis and the beginning of the chronic phase. The proportion of seizure-free rats decreased significantly more slowly in SE+GP1a rats than in SE+Veh rats (Gehan–Breslow–Wilcoxon test, χ²=6.372; p=0.0116; **Fig. 8A**). At the 20th MWM session, 100% of SE+Veh rats had experienced at least one seizure, compared with 65% of SE+GP1a rats. By the end of the two weeks of MWM, SE+Veh rats had almost twice as many induced seizures as SE+GP1a rats (**Fig. 8B**).

**Figure 8.**
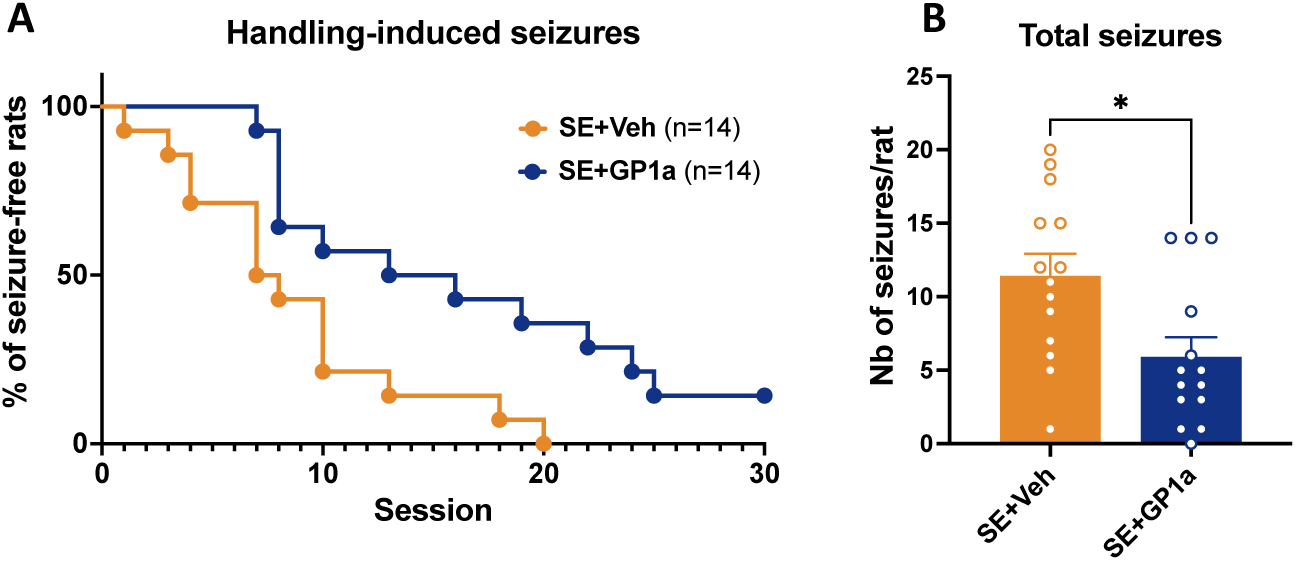
GP1a treatment delays the occurrence of first seizures. Handling-induced seizures were counted manually during the 2 weeks of MWM (3^rd^ and 4^th^ weeks post-SE) in Group 1 rats subjected to SE (n=14/group). **A**. The number of seizure-free rats during MWM sessions decreased significantly slower in GP1a-treated SE rats (Gehan-Breslow-Wilcoxon test, χ²=6.372; p=0.0116). **B**. By the end of the sessions, epileptic rats treated with vehicle had had significantly more induced seizures than GP1a-treated rats (Mann-Whitney: p=0.0087). Results are presented as mean + SEM, *, p<0.05.

## 4. DISCUSSION

Status epilepticus (SE) induced at the juvenile stage elicits a neuroinflammatory response in the hippocampus, characterized by increased expression of both pro- and anti-inflammatory cytokines, robust recruitment of peripheral monocytes, and elevated CB2 expression. We show that selective CB2 activation by the agonist GP1a during epileptogenesis does not blunt the early inflammatory cascade but instead enhances the recruitment of anti-inflammatory monocytes, durably improves cognitive performance and hippocampal synaptic plasticity, reduces anxiety-like behavior, and delays seizure onset. Together, these findings reveal an unexpected mechanism whereby CB2 shapes disease trajectory by modulating microglia–monocyte balance rather than broadly suppressing neuroinflammation.

### Age at SE onset determines inflammatory magnitude and neurodegenerative outcome

Age at SE onset critically influences inflammatory magnitude and long-term pathology. Previous studies demonstrated that juvenile and adult SE differ markedly in inflammatory dynamics and severity, contributing to distinct epilepsy trajectories (Dubé et al., 2001; Scantlebury et al., 2007). In adult models, pro-inflammatory cytokines such as IL-1β, IL-6 and TNFα reach higher levels and remain elevated for prolonged periods, correlating with more pronounced neurodegeneration (Ravizza et al., 2008). In our adult pilocarpine study using identical time points and methods (7h, 1 day, 9 days and 7 weeks post SE), peaks of pro-inflammatory cytokines and persistence of inflammatory markers at 7 weeks were markedly greater (Grabon et al., 2025) than those observed here in the juvenile model. Conversely, in the juvenile brain, inflammation was of lower amplitude, resolved more rapidly and coexisted with sustained anti-inflammatory activation, consistent with reduced vulnerability of immature neurons and restriction of neuronal loss to specific hippocampal subregions (Cilio et al., 2003; De Bruin et al., 2000; Sankar et al., 2000; Scantlebury et al., 2007). These observations suggest that inflammatory magnitude and persistence, rather than mere presence, are critical determinants of neurodegeneration after SE and support therapeutic strategies aiming to modulate, rather than abolish, neuroinflammation in paediatric TLE (Mazarati et al., 2017; Pitkänen et al., 2015; Vezzani et al., 2011).

### Juvenile epileptogenesis is characterized by a monocyte-rich inflammatory program

Within this age specific context, SE was associated with massive recruitment of peripheral monocytes that persisted as brain monocyte-derived macrophages (mo-mΦs) throughout epileptogenesis. Monocyte infiltration after SE is consistent with rodent and human TLE studies showing blood–brain barrier (BBB) leakage and leukocyte recruitment as core features of hippocampal remodeling, and with the identification of MCP-1/CCR2 signaling as a major pathway driving monocyte entry into the epileptic hippocampus (Alemán-Ruiz et al., 2023; Fabene et al., 2008; Morin-Brureau et al., 2018; Ravizza et al., 2008; Reiss et al., 2023; Tian et al., 2017; Varvel et al., 2016; Vezzani et al., 2013; Vinet et al., 2016; Zattoni et al., 2011). Sorted cell analyses revealed that infiltrating monocytes/mo-mΦs display a more pronounced anti-inflammatory profile than microglia and represent the main contributors to increased tissue CB2 transcript levels, whereas CB2 expression in resident microglia remains essentially unchanged (Grabon et al., 2025; Greenhalgh et al., 2016). Following recommendations to prioritize transcript characterization over protein detection with antibodies of disputed specificity (Grabon et al., 2023a), a limitation recently exemplified by the decoupling between robust CB2 PET signals and weaker immunofluorescence (Zeng et al., 2026), we found that CB2 mRNA levels peak at 9 days post-SE. This aligns with the maximal hippocampal PET binding observed during the first week post-SE and supports our choice of a 14-day treatment window (Zeng et al., 2026) using the specific CB2 agonist, GP1a. These findings identify infiltrating monocytes/mo-mΦs as a CB2-positive population particularly amenable to modulation during juvenile epileptogenesis and suggest that CB2 may influence epilepsy outcome primarily through effects on infiltrating myeloid cells.

### CB2 activation reshapes inflammatory composition rather than suppressing acute neuroinflammation

Our data call for a reconsideration of CB2 function in epileptogenesis. CB2 is traditionally viewed as an anti-inflammatory receptor whose activation broadly dampens inflammatory responses and confers neuroprotection in diverse neurological conditions (Grabon et al., 2023; Turcotte et al., 2016). CB2 is predominantly expressed by myeloid cells and its expression is tightly regulated by the inflammatory milieu, with context-dependent regulation reported in sterile versus infectious inflammatory environments (Galiègue et al., 1995; Grabon et al., 2024; Komorowska-Müller and Schmöle, 2020). Contrary to this prevailing view, GP1a treatment during epileptogenesis did not reduce early cytokine induction or classical markers of gliosis, indicating that CB2 does not act upstream of acute microglial activation in this juvenile SE model and contrasting with lesion paradigms where CB2 agonists more directly mitigate microglial activation (Braun et al., 2018; Grabon et al., 2023; Komorowska-Müller and Schmöle, 2020; Zarruk et al., 2012). Instead, CB2 activation markedly increased monocyte recruitment and transiently enhanced the anti-inflammatory response, particularly via IL-4, supporting a model in which CB2 regulates inflammatory composition rather than inflammatory magnitude (Grabon et al., 2025; Greenhalgh et al., 2016; Miller and Stella, 2008).

### Monocyte derived macrophages as context dependent regulators of tissue outcome

The apparent paradox between increased monocyte infiltration and the beneficial effects of GP1a on cognition, anxiety and seizure dynamics can be reconciled by evidence indicating that monocyte-derived macrophages are not intrinsically detrimental and may instead support tissue repair and neuronal survival (Greenhalgh et al., 2016; Kronenberg et al., 2018; Zhang et al., 2019). In a kainate-induced TLE model, depletion of peripheral mononuclear phagocytes reduced F4/80+ macrophage-like cells but also increased granule cell degeneration, suggesting that peripherally derived macrophages contribute to neuronal survival (Zattoni et al., 2011). Conversely, reducing CCR2+ monocyte infiltration in other SE models alleviates inflammation and neuronal damage, highlighting the context dependent nature of monocyte function (Alemán-Ruiz et al., 2023; Tian et al., 2017; Varvel et al., 2016). In our adult pilocarpine study, infiltrating monocytes initially displayed an anti-inflammatory phenotype but later contributed to low grade inflammation once epilepsy was established (Grabon et al., 2025). In the present juvenile model, the GP1a-driven increase in anti-inflammatory mo-mΦs resembles situations in brain injury and neurodegeneration in which monocyte-derived macrophages promote repair and limit secondary damage (Chu et al., 2015; Kronenberg et al., 2018; Shechter et al., 2009; Zhang et al., 2019).

### Methodological considerations strengthen interpretation of myeloid phenotypes

Transcriptional profiles of microglia and monocytes/mo-mΦs were established while minimizing *ex vivo* activation artefacts associated with tissue dissociation and sorting through systematic use of transcription and translation inhibitors at every post-mortem step (Grabon et al., 2025, 2024; Ocañas et al., 2022). This approach strengthens the validity of the *in vivo* inflammatory states attributed to each myeloid population, particularly the early anti-inflammatory phenotype of infiltrating monocytes and the absence of overtly heightened pro-inflammatory activation in chronic mo-mΦs.

### CB2-dependent improvement of cognition and synaptic plasticity indicates disease modification

Beyond immune mechanisms, CB2 activation by GP1a exerted lasting effects on cognition, hippocampal synaptic plasticity and anxiety despite treatment discontinuation before seizure onset, supporting a genuine disease-modifying effect rather than a purely acute anticonvulsant action (Zamarian et al., 2011; Zhao et al., 2014). Our observation of a reduction in handling-induced seizure occurrence following treatment with the GP1a agonist is consistent with a recent study reporting decreased seizure frequency in a pilocarpine mouse model treated with the AM1241 agonist (Cai et al., 2024). Given that hippocampal long-term potentiation (LTP) is a canonical synaptic model of memory substrate (Bliss and Collingridge, 1993), preservation of CA1 LTP by GP1a suggests a direct impact on neuronal mechanisms underlying cognitive improvement. These results are consistent with studies showing that CB2 agonists improve cognitive performance and preserve LTP in neuroinflammatory or neurodegenerative contexts without necessarily normalizing all inflammatory markers (Aso et al., 2013; Martín-Moreno et al., 2012; Wu et al., 2013; Yang et al., 2022).

### Study perspectives and mechanistic considerations

While our findings identify CB2-dependent regulation of monocyte recruitment during juvenile epileptogenesis, the precise contribution of distinct infiltrating monocyte subsets to functional outcomes remains to be established. Future studies combining targeted depletion, lineage tracing or selective chemokine pathway modulation will be instrumental in determining how increased recruitment of these potentially supportive myeloid populations influences neuronal network stability and disease progression. In addition, although transcriptional analyses reveal an anti-inflammatory signature of infiltrating monocytes/mo-mΦs relative to resident microglia, further work will be required to define the functional mechanisms through which these cells modulate hippocampal plasticity and behavioral outcomes. Finally, the restriction of GP1a administration to the epileptogenic phase raises the possibility that early CB2-dependent modulation of myeloid population dynamics may exert long-lasting effects on disease trajectory, a hypothesis that warrants investigation across extended temporal windows. Addressing these questions will refine understanding of CB2-mediated immune regulation and support development of temporally targeted therapeutic strategies.

### Conceptual implications for immune regulation of epileptogenesis

Collectively, these findings redefine the role of CB2 signaling in epileptogenesis by demonstrating that therapeutic benefit arises not from global suppression of neuroinflammation but from selective reorganization of the inflammatory landscape. By promoting recruitment of anti-inflammatory monocyte-derived macrophages without attenuating the initial inflammatory cascade, CB2 activation emerges as a regulator of immune architecture that shapes neuronal network resilience and disease trajectory. This mechanism highlights immune cell composition as a critical determinant of epilepsy outcome and supports a therapeutic framework in which targeted modulation of myeloid population dynamics during epileptogenesis may mitigate both seizure burden and neuropsychiatric comorbidities. More broadly, these results suggest that selective reshaping of inflammatory cell composition represents a promising strategy for disease modification across neuroinflammatory disorders.

## 5. ACKNOWLEDGMENTS

We acknowledge the contribution of SFR Santé Lyon-Est (UAR3453 CNRS, US7 Inserm, UCBL) CyLE cytometry and CIQLE platform facilities, especially Thibault Andrieu and Priscillia Battiston-Montagne for their help with the flow cytometry studies, and Bruno Chapuis for his valuable support with microscopy studies. We gratefully acknowledge Céline Freton (IBCP, Lyon), Pierre Soule, and Cyril Castel (NanoTemper) for their valuable technical assistance with microscale thermophoresis analyses. We thank Nathanaël Leonardi and Jules Dartois for their assistance with behavioral tasks. Nadia Gasmi was granted a PhD fellowship from the Fondation pour la Recherche Médicale. Wanda Grabon was granted a PhD fellowship from France Alzheimer.

## 6. AUTHOR’ S CONTRIBUTION

LB and WG conceived and designed the study. WG, AR, AB, JB, BG, VB, OH and NG participated in data collection. WG, AB, SR and LB analyzed and interpreted the data. WG drafted the manuscript. LB provided critical revisions and approved the final manuscript. All authors read and approved the final manuscript.

## HIGHLIGHTS

- The inflammatory response induced by Li-Pilo-SE at P21 resembles that induced by Pilo-SE at P42, with a lower magnitude.
- Tissue increase in CB2 transcript levels during epileptogenesis is supported by the presence of mo-mΦs.
- CB2 activation during epileptogenesis with the specific agonist GP1a does not alter the inflammatory status of astrocytes, microglia and monocytes/mo-mΦs.
- CB2 activation during epileptogenesis results in a robust increase in the number of protective mo-mΦs.
- Treatment with GP1a improves post-SE recovery, protects cognition, lowers level of anxiety and delays seizure onset.

