## Supplemental data for "CB2 activation during epileptogenesis recruits immunomodulatory monocytes and mitigates cognitive, affective and seizure outcomes"

*****: co-last

### Supplementary material

#### METHODS

##### Experimental design

The experimental design is illustrated in Figure S1. Six distinct groups of animals were similarly subjected to Li-Pilo-SE at P21. Group 1 was used for characterization of the brain inflammatory response following SE and did not receive any treatment. Rats from groups 2 to 6 were treated with the CB2-specific agonist GP1a (Tocris) following SE according to the same protocol: 3mg/kg, in vehicle (5% EtOH, 5%DMSO), at SE+3h, SE+1 day (SE+1D), +2D, +3D, +4D, +7D, +10D, +14D. Eight rats from group 3 were treated according to the same schedule and in the same way in IP, with another CB2 agonist, JWH-133 (Tocris) at a dose of 1.5 mg/kg diluted in the same vehicle. The doses of GP1a and JWH-133 were chosen according to the literature (Braun et al., 2018; Li et al., 2018; Tang et al., 2016; Zarruk et al., 2012).

**Group 6 – Effect of GP1a on hippocampal long-term potentiation.** 16 rats were used in electrophysiological studies to measure the effect of GP1a treatment on Long-Term Potentiation (LTP) measured in fresh hippocampal slices obtained from rats 3-4 weeks after SE (CTRL: n=6; SE: n=5; SE+GP1a: n=5).

Studies conducted to observe the effects of CB2 activation by GP1a on neuroinflammation (Groups 2, 3 and 4) and TLE comorbidities (Groups 5 and 6) are illustrated in Figures S1A and S1B, respectively.

**Group 7 – Effect of electrode implantation on monocyte infiltration**. 5 healthy rats were used to evaluate cellular inflammation at the histological level 1 month following surgical implantation of subdural screw electrodes and intracerebral electrode.

**Animals**

Male Sprague Dawley rats were used in this study (Envigo, France). They were housed under a constant light/dark cycle (12/12h, lights on at 07:00 a.m.), with a room temperature kept at 23 ± 1°C. Animals arrived at 14 days of age (P14) and bred by groups of 10 with a foster mother until weaning. Following weaning, rats were maintained by groups of 2 with *ad libitum* access to food and water in standard cages. The experimental procedures were conducted in compliance with the European Community guidelines for care in animal research and approved by the CELYNE local Ethics Research Committee (protocol APAFIS#12194-2017041813592119 v2). Every effort was made to minimize animal suffering.

**Lithium-pilocarpine Status Epilepticus**

SE was induced by pilocarpine, injected at P21. To prevent peripheral cholinergic side effects, scopolamine methylnitrate (1 mg/kg, s.c.; Sigma-Aldrich) was administered 30 min before pilocarpine hydrochloride (25 mg/kg, i.p.; Sigma-Aldrich). Lithium chloride (127 mg/kg, i.p.; Sigma-Aldrich) was injected 18 hours before scopolamine. After 30 min of continuous behavioral SE, 10 mg/kg diazepam (i.p.; Valium; Roche) was injected, followed 90 min later, by a second injection of 5 mg/kg diazepam to terminate behavioral seizures. The animals were then treated with GP1a or vehicle as described in experimental design. At the end of the SE, the pups were placed back with their foster mother until they regained their weight, and weighed daily.

**Brain collections**

All rats were deeply anesthetized with isoflurane (4%) and administered with a lethal dose of pentobarbital (100 mg/kg; Euthasol). For RT-qPCR analysis (Groups 1 and 2), animals were intracardially perfused with ice-cold saline (30mL/min) to wash out blood from brain vessels. Hippocampi were rapidly microdissected on ice, frozen in liquid nitrogen, and stored at -80°C. For immunochemistry analysis (Groups 1 and 3), animals were intracardially perfused with ice-cold saline to wash out blood from brain vessels and with 4% paraformaldehyde in 0.1 M phosphate buffer (PB) (30mL/min) to fix tissue. For flow cytometry analysis (Group 4), animals were perfused intracardially with ice-cold Dulbecco's PB saline (DPBS) 1X supplemented with Actinomycin D (3µM) Anisomycin (100 µM) and Triptolide (10 µM) i.e. transcription and translation inhibitors. Hippocampi and VLR were quickly dissected and placed in the same buffer until all perfusions were completed.

**Molecular biology**

**RNA extraction –** Tissues (Groups 1 and 2) were crushed in 250 µL of molecular biology grade water (Eurobio) using Tissue-Lyser II (Qiagen) according to the manufacturer's instructions. Total RNAs from brain structures were extracted using Tri-Reagent LS (Molecular Reasearch Center, #TS120). Genomic DNA was removed using Turbo-DNA-free (Ambion, #1907) and total RNAs were purified with RNeasy mini kit (Qiagen, #74104). For sorted cells (Group 4), total RNAs were extracted and purified using the RNeasy Plus Micro Kit (Qiagen, #74034) according to manufacturer's instruction. RNA concentration was determined for each sample on the BioDrop® µLite.

**Reverse transcription and real-time quantitative PCR –** Total tissue RNAs were reverse transcribed to complementary DNA (cDNA) using both oligo dT and random primers with PrimeScript RT Reagent Kit (Takara, #RR037A) according to manufacturer's instructions, in a total volume of 10 µL. In RT reaction, 300 000 copies of a synthetic external non-homologous poly(A) standard messenger RNA (SmRNA; A. Morales and L. Bezin, patent WO2004.092414) were added to normalize the RT step, as previously described (Grabon et al., 2024). cDNA was diluted 1:13 with nuclease free Eurobio water and stored at -20°C until the further use. Each cDNA of interest was amplified from 5 µL of the diluted RT reaction by real-time PCR, using the Rotor-Gene Q thermocycler (Qiagen), the SYBR Green PCR kit (Qiagen, #208052) and oligonucleotide primers (Eurogentec) specific to the targeted cDNA. The sequences of the specific forward and reverse primer pairs were constructed using Primer-BLAST (NCBI). Primers used are listed in **Table S1**. cDNA copy number detected was determined using a calibration curve, and results were expressed as cDNA copy number/µg tot RNA.

Pro-inflammatory and anti-inflammatory indexes were calculated for each series of individuals to be compared using a specific set of genes: IL-1β, IL-6 and TNFα for pro-inflammatory index and IL-4, IL-10 and IL-13 for anti-inflammatory index. For each individual, the number of copies of each transcript has been expressed in percent of the averaged number of copies measured in the whole considered population of individuals. Once each transcript is expressed in percent, an index is calculated by adding the percent of each transcript involved in the composition of the index and expressed in arbitrary units (A.U.).

$$PI = \sum_{i=a}^{n} \frac{cDNA\ copy\ nbr\ for\ transcript\ i}{average\ cDNA\ copy\ nbr\ for\ transcript\ i}$$

with i = IL-1β or IL-6 or TNFα for PI-I and i = IL-4 or IL-10 or IL-13 for AI-I.

**Immunohistology**

**Tissue processing** – 40 µm-thick coronal sections were cut from frozen brains using a cryomicrotome (Leica CM1850; -22°C). Sections were collected in Phosphate Buffer Saline, (PBS 0.1M) and transferred into a cryopreservative solution composed of 19.5 mM $NaH_2PO_4 \cdot 2H_2O$, 19.2 mM NaOH, 30% glycerol and 30% ethyleneglycol and stored at -20°C. Before starting any labeling protocol, free-floating sections were rinsed 3 times with PBS.

**Immunolabelling** – Following permeabilization and saturation with 0.3% Triton 100X and 3% Donkey normal serum, free-floating sections were incubated with primary antibodies. Detection of microglia was performed with goat anti-ionized calcium binding adaptor molecule 1 (Iba1) antibody (1:500, ab5076, Abcam). Infiltrating monocyte-macrophages (mo-MΦ) were labelled with mouse anti-Cluster of differentiation 68 (CD68) antibody (1:1000, MCA341GA, Bio-rad). Finally, astrocytes were double-detected with both mouse and rabbit anti-GFAP (1:1000, G3893, Millipore and 1:1000, ab5804, abcam respectively). For fluorescent dual immunolabeling of Iba-1 and CD68, sections were incubated with an Alexa-Fluor-488-conjugated donkey anti-goat antibody (1:1000, A-11055, Molecular Probes) and an Alexa-Fluor-647-conjugated donkey anti-mouse antibody (1:1000, A-31571, Molecular Probes). For fluorescent dual immunolabeling of GFAP, sections were incubated with Alexa-Fluor-488-conjugated donkey anti-rabbit and anti-mouse antibody (1:1000, A-21206 and A-21202, Molecular Probes), respectively. Nuclei were stained with DAPI (300 nM, Molecular Probes). Sections were mounted on SuperFrost Plus slides and coverglassed with Prolong Diamond Antifade reagent (Molecular Probes)

**Microscopy and analysis** – Whole slices were scanned with a Carl Zeiss Axio Scan.Z1 Digital Slide Scanner (ZEISS) with a X20 lens on a 6 µm stack, using the pilot Zen (ZEISS). Images were then processed on with Fiji software (ImageJ).

**Flow cytometry**

Samples were processed for tissue dissociation and cell sorting immediately following brain collection as quickly as possible. To prevent any artifactual *ex vivo* gene expression changes during brain dissociation and cell sorting procedures, all buffers and solutions used during the process (from animal perfusion to sorted cell flash freezing) were supplemented with a cocktail composed of Actinomycin D (3µM, Tocris #1229/10), Anisomycin (100 µM, Tocris #1290/50) and Triptolide (10 µM, Tocris #3253/10) i.e. transcription and translation inhibitors (Ocañas et al., 2022). All steps were performed on ice or using pre-chilled refrigerated centrifuge set to 4°C with all buffers/solutions pre-chilled before addition to samples to further limit cell activation. Buffer 1 (B1) was composed of DPBS 1X (Thermofisher #14040-117) and inhibitor cocktail. Buffer 2 (B2) was composed of DPBS 1X, BSA 0,5% (Sigma #A2153) and inhibitor cocktail.

**Brain tissue dissociation** – Once collected, tissues were cut in smaller pieces with a scalpel and processed for dissociation using Miltenyi's Adult Brain Dissociation Kit (#130-107-677) according to manufacturer's instruction, running program 37C_ABDK_01. Inhibitor cocktail was added in each reagent. Briefly, samples were added to gentleMACS C Tubes (Miltenyi #130-093-237) with the enzyme mixes and placed in gentleMACS OctoDissociatior with heaters (Miltenyi #130-096-427), running program 37C_ABDK_01. Once program finished, samples were briefly spun before being filtered through 70 µm cell strainer (Thermofisher #11597522). Samples were washed with B1 and spun to pellet cells. To clear cell solution, cell pellets were resuspended and overlaid with appropriate volume of Miltenyi Debris Removal Solution according to manufacturer's protocol. Debris were removed from top layer and solution was diluted with B1 and spun to pellet cells. Cells collected were resuspended in B1 and counted manually (with trypan blue) before magnetic sorting.

**CD11b-positive cells magnetic enrichment** – To increase Fluorescence-Activated Cell Sorting (FACS) yields and efficiency, cell suspensions were first enriched using the Magnetic-Activated Cell Sorting (MACS) technique, magnetically separating CD11b-positive cells (microglia and infiltrating monocytes) for subsequent FACS, from CD11b-negative cells (remaining brain cells, i.e. neurons, astrocytes, oligodendrocytes, endothelial cells, etc.) for direct freezing. In the following steps, cells were suspended in B2. Fc receptors were blocked with anti-CD32 antibody (BD Biosciences #550271). CD11b-positive cells were then enriched using CD11b/c MicroBeads according to manufacturer's instructions (Miltenyi #130-105-634). Briefly, brain cells were incubated with CD11b/c MicroBeads and applied onto MS columns (Miltenyi #130-042-201) in the magnetic field of OctoMACS Separator (Miltenyi #130-042-109). All 8 samples were processed simultaneously. Positive fractions were collected and subjected to the FACS protocol. Unlabelled cells were spun and dry cell pellets were flash frozen and stored at -80°C for further analysis.

**FACS** – Cells were incubated in B2 with anti-CD11b/c PE-conjugated (1:50, BD Biosciences #554862), anti-CD11a BV510-conjugated (1:50, BD Biosciences #744999) and anti-CD45 APCCy7-conjugated (1:50, Biolegend #202216) antibodies. DAPI was used to gate for viable cells. Microglia ($CD11b^{+}CD45^{lo}CD11a^{lo}$) and infiltrating monocytes ($CD11b^{+}CD45^{hi}CD11a^{hi}$) were sorted with BD FACS Aria™ III Cell Sorter (BD Biosciences).

**Behavioral tests**

All behavioral tests were carried out using equipment supplied by Noldus (Netherlands), which provided each test apparatus, camera system, infra-red lighting and tracking software (Ethovision XT video tracking system, Noldus, Netherlands). For all behavioral tests, the running order of animals was determined randomly. In the event of a seizure before or during the task, the rat was returned to its cage for at least 15 min before being retested.

**Elevated Zero-maze –** The elevated zero-maze is a modification of the elevated plus-maze, suggested to have a higher sensitivity to measure anxiety level in rodent (Shepherd et al., 1994). The apparatus consists in an elevated ring 1m in diameter comprising two open portions and two portions protected by dark walls. The rats' movements were recorded for 5 min by video tracking. Infra-red lamps and camera made it possible to measure the rats' trajectories even in the dark areas protected by the walls.

**Water Exploration Test** – As previously described, the WET further allows for the robust measurement of anxiety levels in rodent (Fares et al., 2013). A circular pool (180 cm in diameter) was filled with water maintained at 25°C to a depth of 40 cm and divided into four quadrants of identical size using opaque plastic panels to allow four rats to be recorded simultaneously. The rats' trajectory was recorded for 5 min to measure total distance, average speed and time spent in the virtual central zone considered as anxiogenic.

**Morris Water Maze –** Spatial learning in rats was tested over 5 days using the Morris Water Maze Test (Morris, 1984) in a 180cm circular pool. The water was maintained at 25°C throughout the testing sessions. The tank was divided into 4 virtual quadrants: North (N), East (E), South (S) and West (W). A circular Plexiglas platform (14 cm in diameter) was hidden 1.5 cm below the water surface at a constant position in the north quadrant. The various spatial cues placed on the walls of the room around the pool were kept constant throughout the test. The test was carried out over five consecutive days, with three trials per day separated by a three-hour interval. The first trial on day 1 consisted in positioning the rat on the platform for 60 seconds. For subsequent trials, the starting point of the rats varied in the following order: Day 1, S-S; Day 2, W-W-E; Day 3, E-E-W; Day 4, S-S-E, Day 5, S-W-E. Each test lasted 90 sec. If the animal failed to locate the platform within the allotted time, it was positioned by the experimenter on the platform for 15 sec. A real-time video acquisition system was used to record platform-finding latency, distance and average swimming speed for each trial. The week following MWM spatial learning, retention was assessed during a "probe" session, after removing the platform. Rats' trajectories were recorded for 90 sec to measure latency to platform ex-position and time spent in the target quadrant (N).

**Reversal Morris Water Maze –** To measure spatial learning flexibility, the same experimental paradigm was used immediately afterwards, changing the position of the platform from the N to the S quadrant. On the first day of this second week, rats were positioned for 60 sec on the platform at its new location over two sessions, 3 hours apart. On the following 4 days, the rats were subjected to the same sessions as the MWM week, reversing the position of the starting points. The MWM reversal week was also followed by a 90-second "probe" session without a platform to measure latency to platform ex-position and time spent in both the target quadrant (S) and the previous target (N).

**Strategy analysis during MWM and Reversal MWM tasks** – the Rtrack package was used for the analysis of tracking data, obtained during the MWM and the reversal MWM tasks (https://rupertoverall.net/Rtrack/index.html) with the recommended default parameters. Tracks have been classified in 8 different strategies, illustrated in **Figure S2**.

**Handling-induced seizures** – The number of handling-induced seizures (Kouchi et al., 2022) was strictly reported during all the tests carried out with the Morris pool, i.e. a total of 30 sessions (15 sessions during the MWM test, 13 sessions during the MWM reversal and 2 probe sessions), as these tests were carried out at the end of the epileptogenesis period, i.e. 3-4 weeks following SE.

**Novel Object Recognition task** – The NOR test is widely used test for the investigation into memory alterations (Ennaceur and Delacour, 1988). Rats were first habituated to the empty apparatus, a 1m square arena protected by opaque sidewalls, during a 5 min session. The following day, during the familiarization session, the animals were presented with two identical objects (A and A'), positioned equidistant from the sidewalls. After a 4h delay, the animals were subjected to the testing session, presented with one familiar object (A) and one novel object (B) of similar size, at the same positions. To avoid any possible bias linked to the difference between the two objects, for half of the rats (defined randomly) the familiar object was the one used as new for the other half, and conversely the new object was the one used as familiar for the other half. During both phases of the test, the exploratory behavior of rats was recorded with the videotracking system for a period of 5 min each. Animals were considered to be exploring an object when their nose was detected in the virtual 1 cm zone around the object.

**Electrophysiology**

**Slices preparation** – Transverse hippocampal slices were prepared from P42-49 rats, i.e. 3-4 weeks following SE. Animals were anesthetized using isoflurane and sacrificed by decapitation. Brain was quickly extracted and cooled with ice-cold standard artificial cerebrospinal fluid (ACSF) composed of the following (in mM): 124 NaCl, 5 KCl, 1.25 $Na_2H$-$PO_4$, 2 $MgSO_4$, 2 $CaCl_2$, 26 $NaHCO_3$ and 10 D-Glucose (Sigma-Aldrich). The ACSF was permanently bubbled with 95% $O_2$ and 5% $CO_2$. Hippocampi were dissected and 370 µm-thick transversal slices were prepared using a vibratome (VT1000S; Leica) equipped with a ceramic blade. The slices were then incubated in ACSF at room temperature for at least 1h before transfer to the recording chamber. The ACSF perfused during the recording was supplemented with 100 µM picrotoxin to block $GABA_A$ receptors.

**Electrophysiological recordings** – Whole-cell patch-clamp recordings were obtained from CA1 pyramidal neurons in current clamp mode at -70mV with a patch pipette (3-5 MΩ) containing the following drugs (Sigma-Aldrich): 120 mM potassium gluconate, 20 mM KCl, 0.2 mM EGTA, 2 mM $MgCl_2$, 10 mM HEPES, 4 mM $Na_2ATP$, 0.3 mM Tris-GTP and 14 mM phosphocreatine (pH 7.3, adjusted with KOH). Hippocampal CA1 pyramidal neurons were visualized with a Zeiss Axioskop 2 equipped with a x40 objective, using infrared video microscopy and differential interference contrast optics. The positioning of the patch pipette on the target neuron membrane was performed using a micromanipulator (Luigs & Neumann) under visual control. Whole-cell patch clamp recordings were performed using an Axopatch-200B amplifier (Molecular Devices) at the sampling rate of 10 kHz and filtered at 5 kHz. Data were recorded and analyzed using a Digidata 1440A interface and pClamp 10 software (Molecular Devices). Capillary glass microelectrodes filled with ACSF and connected to an isolator (Iso-Flex, AMPI) were used to stimulate presynaptic axons in the stratum radiatum layer of the hippocampus (120–150 mm away from the soma). Stimulation at 0.05 Hz was used to establish baseline synaptic responses. The stimulation strength was set to evoke excitatory postsynaptic potentials (EPSPs) between 5 and 8 mV. Series resistance (typically 15-25 MΩ) was monitored throughout each experiment; cells with more that 20% change in series resistance were excluded from analysis.

**Long term potentiation** – Once a steady baseline level of EPSPs was recorded for 10 to 20 min, long-term potentiation (LTP) was induced according to the Theta Burst Pairing (TBP) protocol at 5 Hz. A ten-trains sequence separated by 200 ms and composed of 5 pulses of 1 ms at 100 Hz coupled to backward action potentials (AP) is repeated 3 times at 10 seconds intervals. 0,05 Hz stimulations were resumed immediately after tetanization, and the amplitude of the EPSP recorded for 40 min. LTP is expressed as a percentage of increase in responses, relative to the baseline level previously recorded. Electrophysiological data were analyzed using pClamp 10 and Igor pro software (WaveMetrics).

**Stereotactic surgery**

General anesthesia was induced by administration of a ketamine (80 mg/kg) and xylazine (10 mg/kg) mix, i.p and local analgesia was induced by subcutaneous injection of lidocaine (5 mg/kg). Intracerebral electrode (stainless two-twisted teflon-coated wires, 20 mm length, #8IE36332TWXE, Bilaney Consultants) was positioned in the right basolateral amygdala (coordinates relative to bregma in mm, according to Paxinos and Watson rat brain atlas, rostrocaudal: -2.8; mediolateral: +4.8; dorsoventral: -8.5). Subdural screw electrode (stainless steel screw, Ø1.6 mm, #E363/96/1.6/SPC, Bilaney Consultants) was positioned over the right frontal cortex. The leads from both electrodes were inserted into a plastic pedestral (#MS363 pedestral 2298 6 PIN, Plastic Ones) and fixed using dental acrylic (Paladur®). Metacam (1 mg/kg, per os) was administered after surgery and for the following two days.

**Microscale thermophoresis**

To check the affinity of GP1a for CB2, binding experiments were carried out by microscale thermophoresis (MST). Mouse mCB2-HIS-Tag was expressed and assembled into nanodiscs (Creative Biomart). MST enables to track temperature-induced changes in the fluorescence of a fluorochrome-tagged protein. When a ligand binds to a target, its MST signal is modified in a ligand concentration-dependent manner (Wienken et al., 2010). Transition of the signal from the saturated-protein state (high ligand concentration) to the free-protein state (low ligand concentration) was monitored to determine the dissociation constant ($K_d$). The His-Tag domain was labelled using the Monolith Protein Labeling Kit RED-NHS according to the manufacturer's instructions. mCB2-HIS-Tag (5 nM) was incubated 20 min with GP1a at 16 different concentrations obtained by serial dilution (from 62.5 nM to 1.9 pM) in buffer DPBS 0.01% pluronic. Samples were loaded into glass capillaries (Monolith NT Capillaries, Nanotemper Technologies), and thermophoresis analysis was performed using a Monolith NT.115 Series instrument (Nanotemper Technologies, excitation power 100%). Signal quality was monitored by the NanoTemper Monolith device to detect possible ligand auto-fluorescence, precipitation, aggregation, or ligand-induced changes in the photobleaching rate. Experiments were conducted as triplicates and treated with the MO.Affinity Analysis software (NanoTemper).

**Statistical Analysis**

Statistical analyses were performed using Prim 10.0 software (GraphPad, USA). Results are presented as mean + SEM (standard error of the mean). Differences with a p-value<0.05 ($p<0.05$) were considered to be statistically significant. The Shapiro–Wilk test and quantile–quantile plot were used to assess normal distribution of the data. For the data with normal distribution, the statistical significance was assessed by t-test, one-way or two-way ANOVA analysis, followed with Tukey's post-hoc test for multiple comparisons. For non-normal distribution data, the statistical significance was assessed by Kruskal-Wallis' test, followed with Dunn's post-hoc test for multiple comparisons or Mann-Whitney test for two groups comparisons. Simple linear regression was used to assess the association between two variables. The transcript values of one of the rats in the SE+Veh 10h group were considered outlier (> mean + 2 times standard deviation or < mean - 2 times standard deviation) for 4 of the 7 genes presented here and was therefore excluded from the study. Details of statistic tests for each figure are presented in supplementary data (**Table S2**).

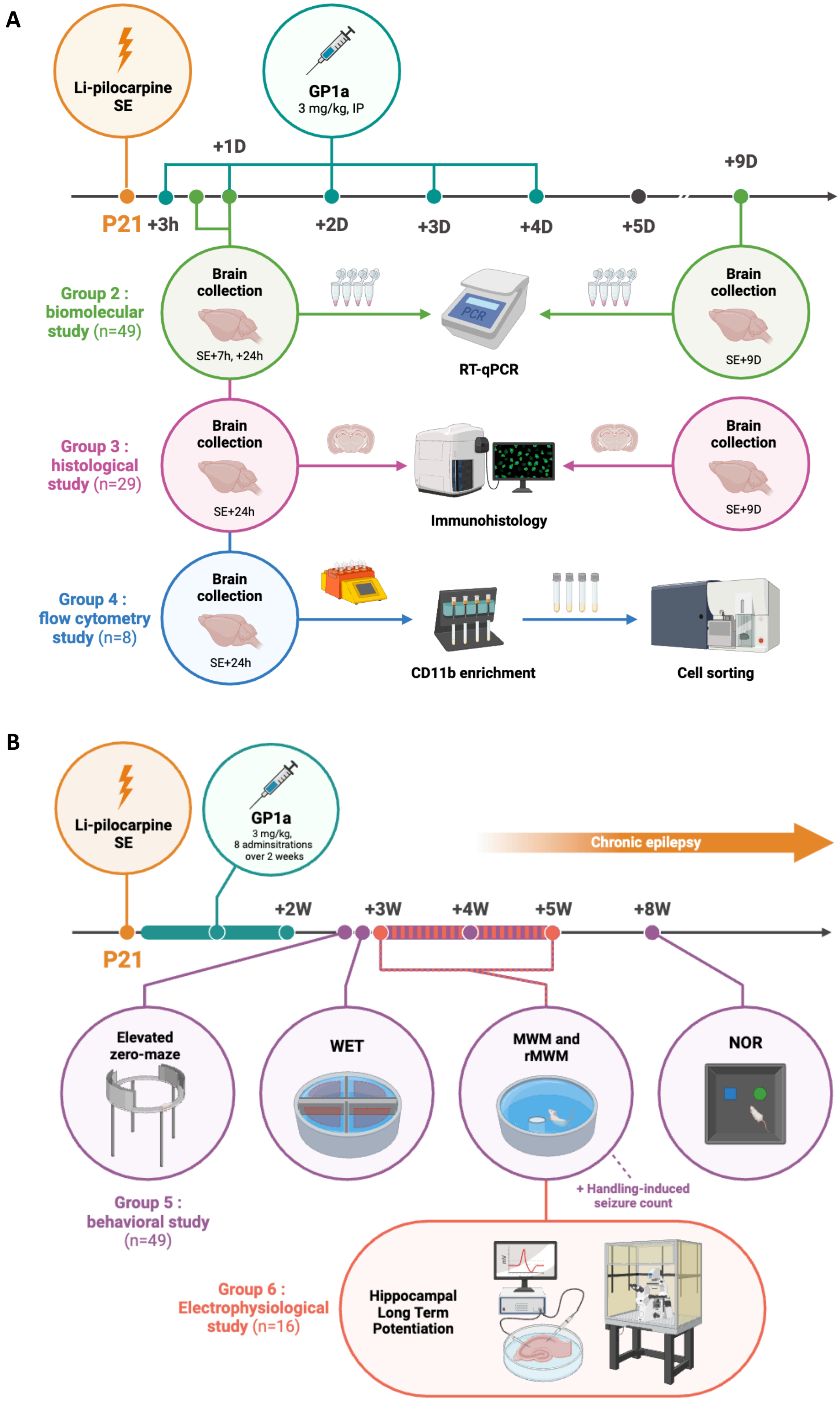


**Figure S1 – Experimental design.** Studies conducted to measure the effects of CB2 activation by GP1a on **A.** SE-induced neuroinflammation (Groups 2, 3 and 4) and **B.** TLE comorbidities (Groups 5 and 6).

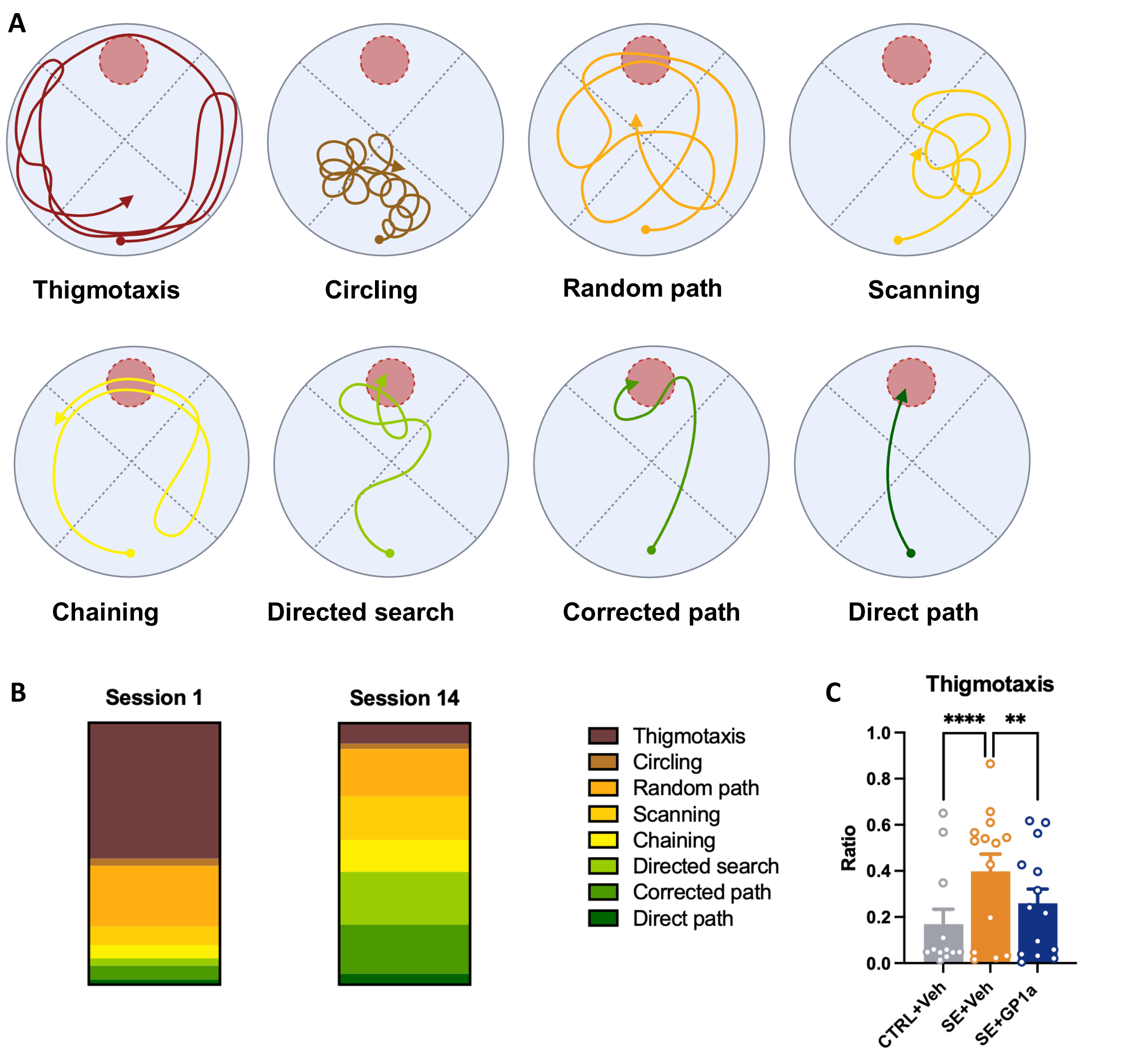


**Figure S2 – Strategy analysis during MWM and Reversal MWM tasks. A.** Representation of the eight strategies, as defined in Rtrack (https://rupertoverall.net/Rtrack/index.html), that can be grouped into three functional categories: non-oriented strategies (thigmotaxis, circling and random path), which reflect non-oriented exploration and/or wall-associated behavior; procedural strategies (scanning and chaining), based on repetitive or egocentric search patterns; and allocentric strategies (directed search, corrected path and direct path), which reflect increasingly efficient spatial navigation toward the platform using distal environmental cues. **B**. Evolution of the average distribution of swim-search strategies in control vehicle-trated rats (CTRL+Veh; n=14) between the first and the last Morris Water Maze (MWM) sessions (sessions 1 and 14), illustrating the shift from predominantly non-goal-oriented to more efficient spatial search strategies during task acquisition. **C**. Proportion of trials classified as thigmotaxis strategy during Probe Trial 2 in CTRL+Veh (n=12), in SE+Veh (n=14) and in SE+GP1a (n=14). Data are shown as individual rats with mean + SEM. Details of statistic tests are provided in supplementary table 2. **, $p<0.01$; ****, $p<0.0001$.

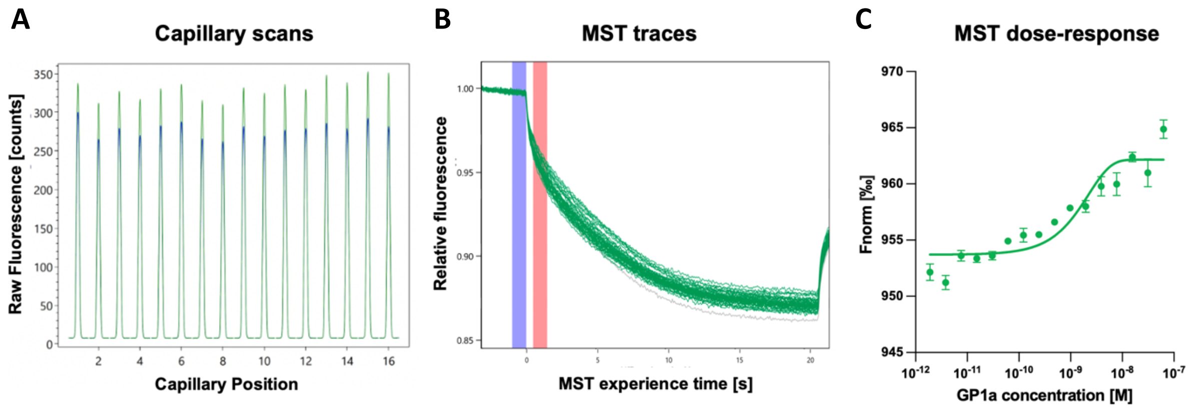


**Figure S3 – GP1a strongly binds to CB2.** MST assays were used to assess GP1a binding capacity and affinity for CB2 *in vitro* using mCB2-HIS-Tag expressed and assembled into nanodiscs. **A.** Visualization of capillary scans, to check for the absence of adsorption and ligand-induced fluorescence change. **B.** Visualization of cumulative MST traces, to check for the absence of protein aggregation and ligand-induced photobleaching. **C.** MST dose-response curve of GP1a: data are presented as mean ± SEM. $K_d$ for GP1a was estimated at 1.19 ± 0.52 nM by MO.Affinity Analysis software.

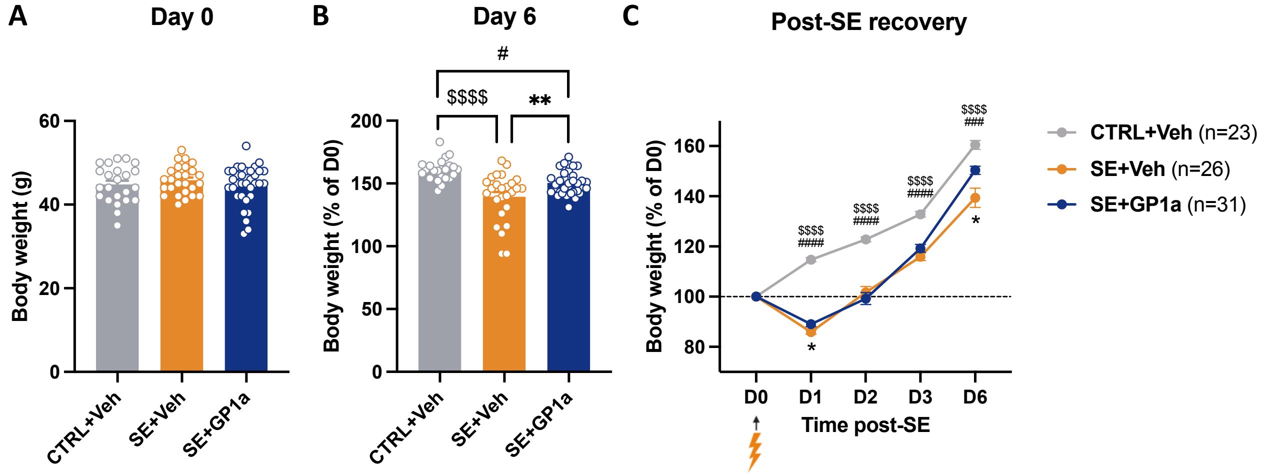


**Figure S4 – GP1a-treated rats recovered faster following SE**. Body weight of rats was monitored after SE and compared to rats that were not subjected to SE to evaluate recovery, in all rats included in this study that were followed for 6 days or more after SE (Group 2 for tissue RT-qPCR studies; Group 3 for histological studies; Group 5 for behavioral studies and Group 6 for electrophysiological studies; total: CTRL+Veh, n=23; SE+Veh, n=26; SE+GP1a, n=31. See groups details in Method section). **A**. Rats were weighted 1h prior to SE induction. These measurements served as a reference for the determination of changes in the body weight during the follow-up. **B-C**. Body weight was expressed as percent of day 0 following SE. Details of statistical tests are presented in Table S2. $: CTRL+Veh vs SE+Veh; #: CTRL+Veh vs SE+GP1a; *: SE+Veh vs SE+GP1a. */$/#, p<0.05; **/$$/##, p<0.01; ***/$$$/####, p<0.001; ****/$$$$/####, p<0.0001.

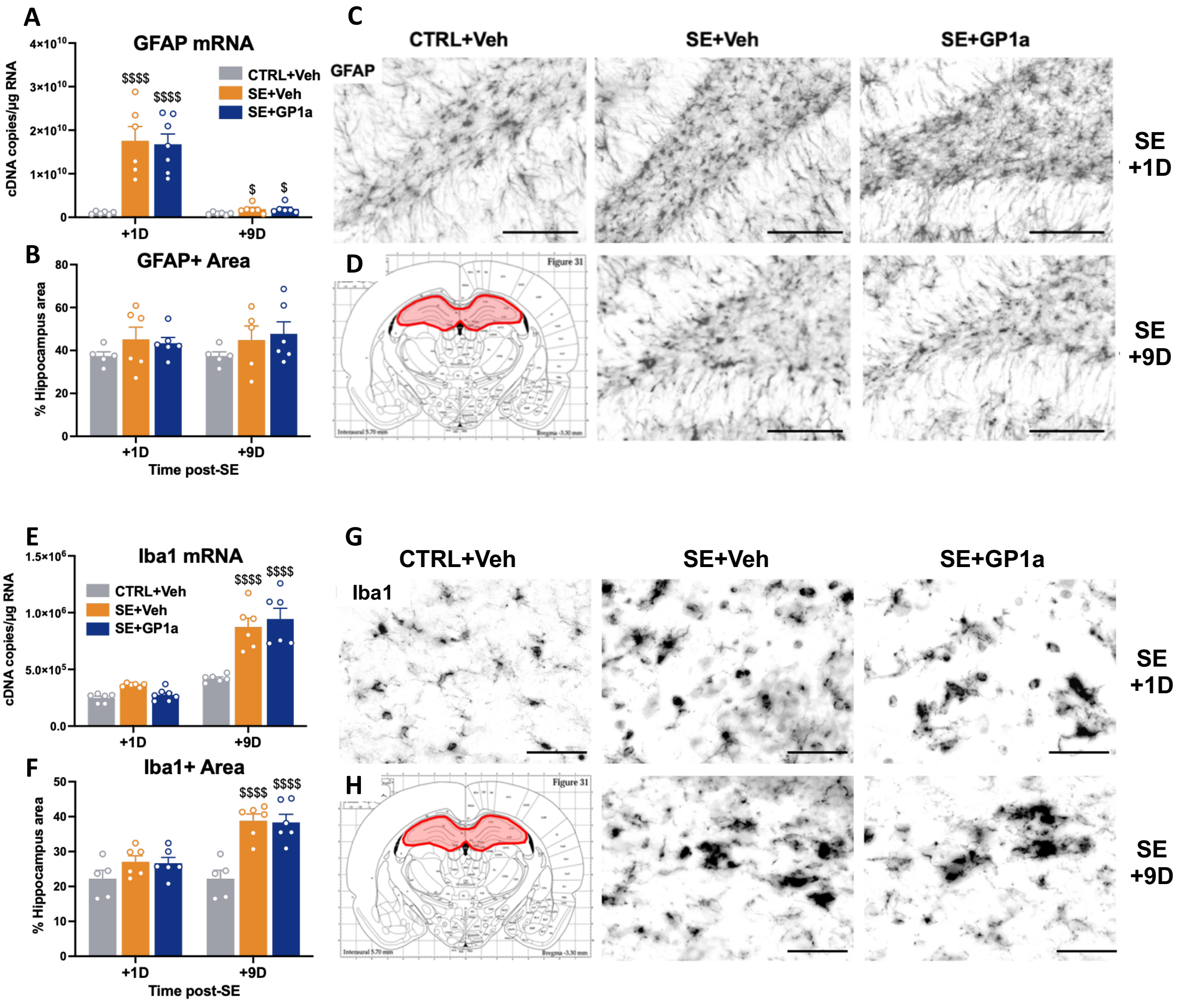


**Figure S5 – GP1a treatment does not influence astrocyte and microglia activation in the hippocampus. A-D. Astrogl iosis. A.** SE-induced hippocampal astrogliosis was quantified by measuring GFAP transcript levels by RT-qPCR in the hippocampus 1D and 9D following SE collected from Vehicle-treated (SE+Veh, n=5-6/time point) and GP1a-treated SE rats (SE+GP1a, n=6/time point) and compared with healthy controls (CTRL+Veh, n=5), in Group 2 rats. **B-D**. Hippocampal astrogliosis was further evaluated by immunohistochemistry, by quantifying GFAP-positive area in the hippocampus 1D and 9D following SE in Vehicle-treated (SE+Veh, n=6/time point) and GP1a-treated SE rats (SE+GP1a, n=6/time point) and compared with healthy controls (CTRL+Veh, n=5), in Group 3 rats. **B**. Quantification of GFAP-positive area, expressed as percent of total hippocampus surface area. **C**. Representative histological images of GFAP immunodetection (anti-GFAP antibodies: Millipore G3893 and abcam ab5804), acquired with slide scanner in the dentate gyrus hilus (x20 objective scale bars: 100µm). **D**. In red: delimitation of the region where the quantifications were carried out, i.e. dorsal hippocampus at bregma -3.30mm). **E-K. Microgliosis. E.** Microglia activation was estimated by measuring Iba1 transcript levels. **F-H**. Hippocampal microglia activation was further evaluated by immunohistochemistry. **G**. Quantification of Iba1-positive area, expressed as percent of total hippocampus surface area. **G**. Representative histological images of Iba1 immunodetection (anti-Iba1 antibody: abcam ab5076), acquired with slide scanner in the dentate gyrus hilus (x20 objective scale bars: 50µm). **H**. In red: delimitation of the region where the quantifications were carried out, as for astrocytes. Details of statistic tests are presented in supplementary data. Results are presented as mean + SEM. $: vs CTRL+Veh; *: SE+Veh vs SE+GP1a. */$, p<0.05; **/$$, p<0.01; ***/$$$, p<0.001; ****/$$$$, p<0.0001.

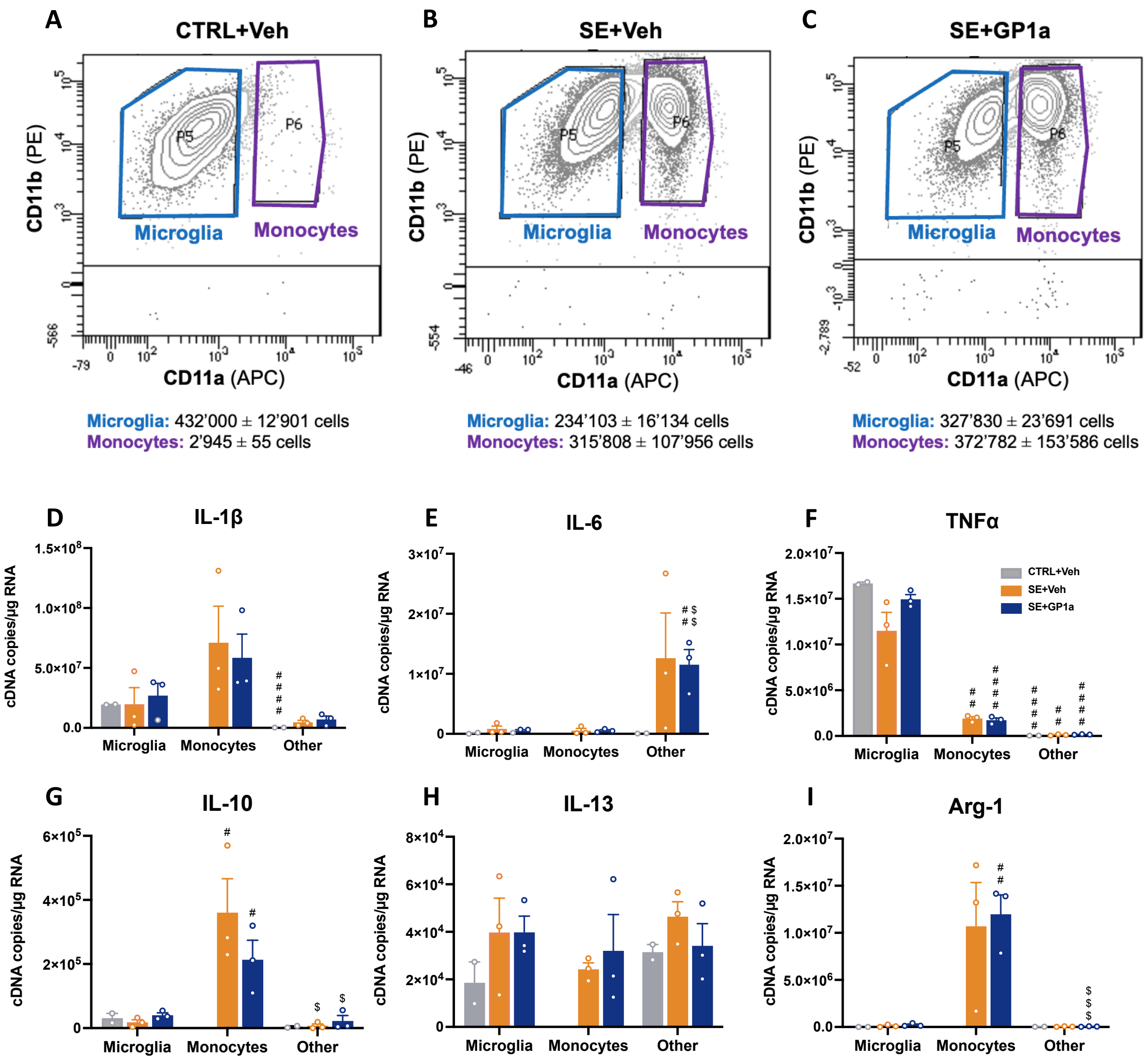


**Figure S6 – Infiltrating monocytes are less inflammatory than microglia 1D following SE. A-C.** Microglia (CD11b$^{+}$CD45$^{lo}$CD11a$^{lo}$) and monocytes (CD11b$^{+}$CD45$^{hi}$CD11a$^{hi}$) were sorted by FACS following CD11b MACS enrichment from hippocampus and VLR collected in Group 4 rats 1D following SE (CTRL+Veh, n=2; SE+Veh, n=3; SE+GP1a, n=3). CD11b-negative fractions were also collected following MACS enrichment. **D-I**. Transcripts of pro-inflammatory (**D.** IL-1β, **E.** TNF-α and **F.** IL-6) and anti-inflammatory (**G.** IL-10, **H.** IL-13 and **I.** Arg-1) associated genes were quantified on sorted cell populations by RT-qPCR and expressed as cDNA copies per microgram of total RNA. Details of statistic tests are presented in supplementary data. Results are presented as mean + SEM. Inter-cell type differences: #, vs respective microglia; $: vs respective mo-mΦs. #, p<0.05; ##, p<0.01; ###, p<0.001; ####, p<0.0001.

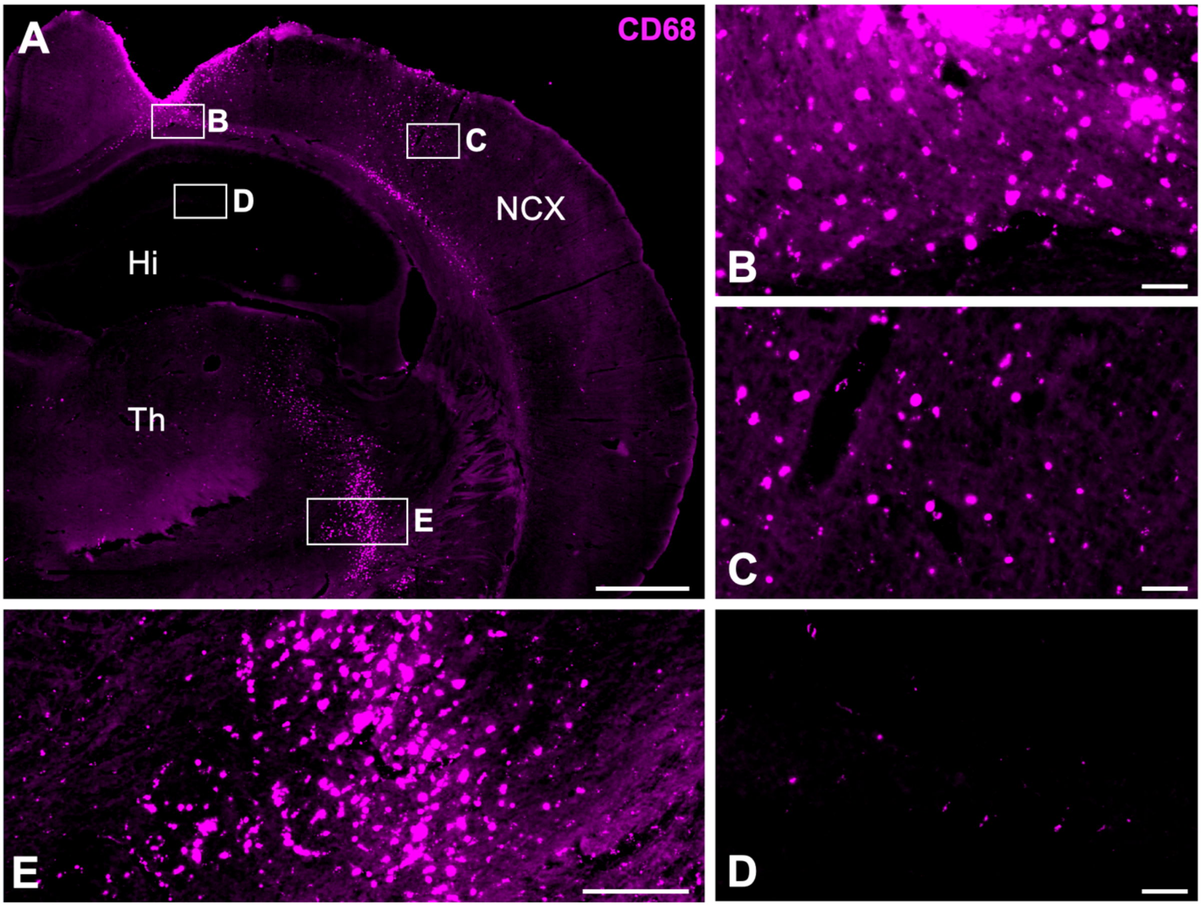


**Figure S7 – Scalp and intracerebral electrode implantation results in monocyte infiltration in the brain of healthy rats.** Healthy rats were implanted with subdural screw electrode (above the neocortex) and intracerebral (in the baso-lateral amygdala) recording electrodes to observe subsequent monocyte infiltration and presence in brain tissue one month post-surgery. **A**. Infiltrating monocytes were immunolabeled with anti-CD68 antibody (Biorad MCA341GA). **B-E**. Magnifications of image A in the neocortex (B, C), the dorsal hippocampus (D) and the thalamus (E). Scale bars: A, 1000µm; B-D, 50µm, E, 200µm. **Abbreviations**: Hi, hippocampus; NCX, neocortex; Th, thalamus.

**Table S1. Sequences of primer pairs used for qPCR** (*Rattus Norvegicus*)

**Abbreviations**: Arg1, arginase 1; CB2, cannabinoid receptor type 2; CD68, cluster of differentiation 68; GFAP,

| Target cDNA | Forward primer sequence | Reverse primer sequence | GenBank Reference |
|---|---|---|---|
| **Arg1** | 5' TCC AAG CCA AAG CCC ATA GAG 3' | 3' CTT TGT ATG TTA CAC TCT CTG 5' | NM_017134.3 |
| **CB2** | 5' TGA CCG CTG TTG ACC GAT AC 3' | 3' CGA GAG GAC CCA CAT GAC AC 5' | NM_020543 (*) |
| **CD68** | 5' CTT TCT CCA GCA ATT CAC CTG 3' | 3' ACT GGC GCA AGA GAA GCA 5' | NM_001031638.1 |
| **GFAP** | 5' ACA TCG AGA TCG CCA CCT AC 3' | 3' GGA TCT GGA GGT TGG AGA AA 5' | NM_017009.2 |
| **Iba1** | 5' CCA GCG TCT GAG CTA TG 3' | 3' CCA GCA TTC GCT TCA AGG AC 5' | NM_01716.3 |
| **IL-10** | 5' TTG AAC CAC CCG GCA TCT AC 3' | 3' CCA AGG AGT TGC TCC CGT TA 5' | NM_012854.2 |
| **IL-13** | 5' AGT CCT GGC TCT CGC TTG 3' | 3' GAT GTG GAT CTC CGC ACT G 5' | NM_053828.1 |
| **IL-1β** | 5' TGT GAT GAA AGA CGG CAC AC 3' | 3' CTT CTT CTT TGG GTA TTG TTT GG 5' | NM_031512.2 |
| **IL-4** | 5' GTA GAG GTG TCA GCG GTC TG 3' | 3' TTC AGT GTT GTG AGC GTG GA 5' | NM_201270.1 |
| **IL-6** | 5' CCC TTC AGG AAC AGC TAT GAA 3' | 3' ACA ACA TCA GTC CCA AGA AGG 5' | NM_012589.1 |
| **MCP1** | 5' CGG CTG GAG AAC TAC AAG AGA 3' | 3' TCT CTT GAG CTT GGT GAC AAA TA 5' | NM_57441.1 |
| **TNFα** | 5' TGA ACT TCG GGG TGA TCG 3' | 3' GGG CTT GTC ACT CGA GTT TT 5' | NM_012675.3 |
| **SmRNA** | 5' CGG GAC AAG AAG GTG GAA G 3' | 5' AGT CTG CAG TGA GTC GAA GAA A 3' | WO2004.092414 |

GFAP, Glial fibrillary acidic protein; Iba1, ionized calcium-binding adapter molecule 1; IL, interleukin; MCP1, monocyte chemoattractant protein 1; TNFα, tumor necrosis factor α; SmRNA, standard messenger RNA.
* The CB2 primers amplify a sequence shared by several transcripts. As indicated in the table, NM_020543 is the identified transcript, although the same sequence is also present in NM_001164142, NM_001164143, and NM_001416630.

**Table S2 – Details of statistic tests.** Normality of data distribution was tested using Shapiro–Wilk test and quantile–quantile plots ("Normality?" column) and the statistic test was chosen accordingly for each group.

**Figure 1**

| | | | p-value | | | | Normality ? | Test | Post-hoc |
|---|---|---|---|---|---|---|---|---|---|
| | | | vs. CTRL (*) | | vs. SE+7h ($) | | | | |
| 1A | IL-1β | SE+7h | <0.0001 | **** | | | No | Krukall Wallis | Dunn's |
| | | SE+1D | 0.0289 | * | 0.2737 | ns | | | |
| | | SE+9D | 0.5579 | ns | 0.0029 | $$ | | | |
| 1B | IL-6 | SE+7h | 0.0065 | ** | | | Yes | One-way ANOVA | Tukey's |
| | | SE+1D | 0.9998 | ns | 0.0025 | $$ | | | |
| | | SE+9D | >0.9999 | ns | 0.0012 | $$ | | | |
| 1C | TNFα | SE+7h | <0.0001 | **** | | | Yes | One-way ANOVA | Tukey's |
| | | SE+1D | 0.9893 | ns | <0.0001 | $$$$ | | | |
| | | SE+9D | 0.9893 | ns | <0.0001 | $$$$ | | | |
| 1D | Pro-inflam index | SE+7h | 0.0088 | ** | | | No | Krukall Wallis | Dunn's |
| | | SE+1D | >0.9999 | ns | 0.1575 | ns | | | |
| | | SE+9D | >0.9999 | ns | 0.0002 | $$$ | | | |
| 1E | IL-4 | SE+7h | 0.1072 | ns | | | Yes | One-way ANOVA | Tukey's |
| | | SE+1D | 0.0420 | * | 0.9741 | ns | | | |
| | | SE+9D | >0.9999 | ns | 0.0375 | $ | | | |
| 1F | IL-10 | SE+7h | <0.0001 | **** | | | Yes | One-way ANOVA | Tukey's |
| | | SE+1D | <0.0001 | **** | 0.7089 | ns | | | |
| | | SE+9D | 0.8345 | ns | <0.0001 | $$$$ | | | |
| 1G | IL-13 | SE+7h | 0.0011 | ** | | | Yes | One-way ANOVA | Tukey's |
| | | SE+1D | 0.0215 | * | 0.4853 | ns | | | |
| | | SE+9D | 0.9961 | ns | <0.0001 | $$$$ | | | |
| 1H | Anti-inflam index | SE+7h | 0.0013 | ** | | | Yes | One-way ANOVA | Tukey's |
| | | SE+1D | 0.0017 | ** | 0.9208 | ns | | | |
| | | SE+9D | 0.9208 | ns | 0.0001 | $$$ | | | |

**Figure 2**

| | | | p-value | | | | Normality ? | Test | Post-hoc |
|---|---|---|---|---|---|---|---|---|---|
| | | | vs. CTRL (*) | | vs. SE+7h ($) | | | | |
| 2A | GFAP | SE+7h | 0.4863 | ns | | | Yes | One-way ANOVA | Tukey's |
| | | SE+1D | <0.0001 | **** | 0.0001 | $$$ | | | |
| | | SE+9D | 0.0099 | ** | 0.1748 | ns | | | |
| 2B | Iba1 | SE+7h | 0.0230 | * | | | Yes | One-way ANOVA | Tukey's |
| | | SE+1D | 0.9351 | ns | 0.0397 | $ | | | |
| | | SE+9D | 0.3856 | ns | <0.0001 | $$$$ | | | |
| 2C | MCP-1 | SE+7h | <0.0001 | **** | | | Yes | One-way ANOVA | Tukey's |
| | | SE+1D | 0.0002 | *** | <0.0001 | $$$$ | | | |
| | | SE+9D | >0.9999 | ns | <0.0001 | $$$$ | | | |
| 2D | CD68 | SE+7h | >0.9999 | ns | | | No | Krukall Wallis | Dunn's |
| | | SE+1D | 0.0347 | * | 0.1867 | ns | | | |
| | | SE+9D | 0.0003 | *** | 0.0024 | $$ | | | |

**Figure 3**

| | | p-value | | Normality ? | Test | Post-hoc |
|---|---|---|---|---|---|---|
| | | Microglia vs. mo-mΦs | | | | |
| 3D | IL-1β | 0.2004 | ns | Yes | Unpaired t-test | |
| | TNFα | 0.0091 | ** | Yes | Unpaired t-test | |
| | IL-6 | 0.4000 | ns | No | Mann-Whitney | |
| | IL-10 | 0.0321 | * | Yes | Unpaired t-test | |
| | IL-13 | 0.3518 | ns | Yes | Unpaired t-test | |
| | Arg1 | 0.0062 | ** | Yes | Unpaired t-test | |

| | | | p-value | | | | Normality ? | Test | Post-hoc |
|---|---|---|---|---|---|---|---|---|---|
| | | | vs. CTRL (*) | | vs. SE+7h ($) | | | | |
| 3E | CB2 (Tissue) | SE+7h | 0.7134 | ns | | | Yes | One-way ANOVA | Tukey's |
| | | SE+1D | 0.1238 | ns | 0.0500 | $ | | | |
| | | SE+9D | 0.0009 | *** | <0.0001 | $$$$ | | | |
| | | | vs. CTRL Microglia (*) | | vs. SE Microglia (*) | | Normality ? | Test | Post-hoc |
| 3F | CB2 (FACS) | Microglia | 0.8054 | ns | | | Yes | One-way ANOVA | Tukey's |
| | | mo-mΦs | 0.0415 | * | 0.0905 | ns | | | |
| | | Other | 0.0004 | *** | 0.0005 | *** | | | |

**Figure 4**

| | | | p-value | | | | | | Normality ? | Test | Post-hoc |
|---|---|---|---|---|---|---|---|---|---|---|---|
| | | | CTRL+Veh vs. SE+Veh ($) | | CTRL+Veh vs. SE+GP1a ($) | | SE+Veh vs. SE+GP1a (*) | | | | |
| **4A** | **CB2** | **SE+10h** | 0,3879 | ns | 0.0374 | $ | 0.0035 | ** | Yes | | Tukey's |
| | | **SE+1D** | <0.0001 | $$$$ | <0.0001 | $$$$ | 0.9794 | ns | Yes | One-way ANOVA | Tukey's |
| | | **SE+9D** | 0.0018 | $$ | 0.0005 | $$$ | 0.7688 | ns | Yes | | Tukey's |
| **4B** | **Pro-inflam index** | **SE+10h** | 0.0127 | $ | 0.0025 | $$ | 0.7921 | ns | Yes | | Tukey's |
| | | **SE+1D** | 0.3223 | ns | 0.0604 | ns | 0.6259 | ns | Yes | One-way ANOVA | Tukey's |
| | | **SE+9D** | 0.0088 | $$ | 0.0074 | $$ | 0.9959 | ns | Yes | | Tukey's |
| **4C** | **Anti-inflam index** | **SE+10h** | 0.0135 | $ | <0.0001 | $$$$ | 0.0125 | * | Yes | | Tukey's |
| | | **SE+1D** | 0.1160 | ns | 0.1370 | ns | 0.9832 | ns | Yes | One-way ANOVA | Tukey's |
| | | **SE+9D** | 0.7680 | ns | 0.5360 | ns | 0.2078 | ns | Yes | | Tukey's |
| **4D** | **IL-1β** | **SE+10h** | 0.0001 | $$$ | 0.0002 | $$$ | 0.9936 | ns | Yes | | Tukey's |
| | | **SE+1D** | 0.1358 | ns | 0.2634 | ns | 0.8791 | ns | Yes | One-way ANOVA | Tukey's |
| | | **SE+9D** | <0.0001 | $$$$ | <0.0001 | $$$$ | 0.9977 | ns | Yes | | Tukey's |
| **4E** | **IL-6** | **SE+10h** | 0.0234 | $ | <0.0001 | $$$$ | 0.0071 | ** | Yes | One-way ANOVA | Tukey's |
| | | **SE+1D** | 0.0267 | $ | 0.0028 | $$ | >0.9999 | ns | No | Kruskall Wallis | Dunn's |
| | | **SE+9D** | 0.5286 | ns | 0.3604 | ns | 0.9483 | ns | Yes | One-way ANOVA | Tukey's |
| **4F** | **TNFα** | **SE+10h** | 0.0393 | $ | 0.1576 | ns | 0.6652 | ns | Yes | One-way ANOVA | Tukey's |
| | | **SE+1D** | 0.1943 | ns | 0.4920 | ns | >0.9999 | ns | No | Kruskall Wallis | Dunn's |
| | | **SE+9D** | 0.3702 | ns | 0.3097 | ns | 0.9908 | ns | Yes | One-way ANOVA | Tukey's |
| **4G** | **IL-4** | **SE+10h** | 0.0057 | $$ | <0.0001 | $$$$ | 0.0078 | ** | Yes | One-way ANOVA | Tukey's |
| | | **SE+1D** | 0.0310 | $ | 0.0024 | $$ | >0.9999 | ns | No | Kruskall Wallis | Dunn's |
| | | **SE+9D** | >0.9999 | ns | >0.9999 | ns | >0.9999 | ns | Yes | One-way ANOVA | Tukey's |
| **4H** | **IL-10** | **SE+10h** | 0.5120 | ns | 0.0107 | $ | 0.1180 | ns | Yes | | Tukey's |
| | | **SE+1D** | 0.1666 | ns | 0.2737 | ns | 0.9214 | ns | Yes | One-way ANOVA | Tukey's |
| | | **SE+9D** | 0.6811 | ns | 0.0815 | ns | 0.3243 | ns | Yes | | Tukey's |
| **4I** | **IL-13** | **SE+10h** | 0.0039 | $$ | 0.0125 | $ | 0.7366 | ns | Yes | One-way ANOVA | Tukey's |
| | | **SE+1D** | 0.2714 | ns | >0.9999 | ns | >0.9999 | ns | No | Kruskall Wallis | Dunn's |
| | | **SE+9D** | 0.1918 | ns | 0.5455 | ns | 0.7295 | ns | Yes | One-way ANOVA | Tukey's |

#### Figure 5

| | | | p-value | | | | | | Normality ? | Test | Post-hoc |
|---|---|---|---|---|---|---|---|---|---|---|---|
| | | | CTRL+Veh vs. SE+Veh ($) | | CTRL+Veh vs. SE+GP1a ($) | | SE+Veh vs. SE+GP1a (*) | | | | |
| **5A** | **MCP-1 mRNA** | **SE+10h** | 0.0017 | $$ | <0.0001 | $$$$ | 0.0118 | * | Yes | One-way ANOVA | Tukey's |
| | | **SE+1D** | <0.0001 | $$$$ | <0.0001 | $$$$ | 0.6828 | ns | Yes | One-way ANOVA | Tukey's |
| | | **SE+9D** | 0.1980 | ns | >0.9999 | ns | >0.9999 | ns | No | Kruskall Wallis | Dunn's |
| **5B** | **CD68+ Area** | **SE+1D** | 0.0096 | $$ | 0.0264 | $ | >0.9999 | ns | No | Kruskall Wallis | Dunn's |
| | | **SE+9D** | 0.2286 | ns | 0.0002 | $$$ | 0.0036 | ** | Yes | One-way ANOVA | Tukey's |
| **5C** | **Density of CD68+ cells** | **SE+1D** | 0.0109 | $ | 0.0028 | $$ | 0.8407 | ns | Yes | Two-way ANOVA | Tukey's |
| | | **SE+9D** | 0.0011 | $$ | <0.0001 | $$$$ | 0.0456 | * | | | |
| | | | **SE+Veh vs. SE+GP1a (*)** | | | **SE+Veh vs. SE+JWH-133 (*)** | | | **Normality ?** | **Test** | **Post-hoc** |
| **6G** | **CD68+ Area** | | 0.0070 | | ** | 0.0247 | | * | Yes | Unpaired t-test | |
| **6H** | **Density of CD68+ cells** | | 0.0083 | | ** | 0.0316 | | * | Yes | Unpaired t-test | |

#### Figure 6

| | | | p-value | | | | | | Normality ? | Test | Post-hoc |
|---|---|---|---|---|---|---|---|---|---|---|---|
| | | | CTRL+Veh vs. SE+Veh ($) | | CTRL+Veh vs. SE+GP1a (#) | | SE+Veh vs. SE+GP1a (*) | | | | |
| **6B** | **Day 1** | | 0.6901 | ns | 0.5920 | ns | 0.5897 | ns | Yes | Two-way repeated measures ANOVA | Tukey's |
| | **Day 2** | | 0.4404 | ns | 0.7914 | ns | 0.8892 | ns | | | |
| | **Day 3** | | 0.3582 | ns | 0.1402 | ns | 0.7807 | ns | | | |
| | **Day 4** | | 0.0478 | $ | 0.0242 | # | 0.9603 | ns | | | |
| | **Day 5** | | 0.0656 | ns | 0.2097 | ns | 0.6817 | ns | | | |
| | | | **CTRL+Veh** | | **SE+Veh** | | **SE+GP1a** | | | | |
| | **D1 vs. D4** | | 0.0067 | $$ | 0.0174 | # | 0.0126 | * | | | |
| | **D1 vs. D5** | | 0.0003 | $$$ | 0.0183 | # | 0.0003 | *** | | | |
| | | | **CTRL+Veh vs. SE+Veh ($)** | | **CTRL+Veh vs. SE+GP1a (#)** | | **SE+Veh vs. SE+GP1a (*)** | | **Normality ?** | **Test** | **Post-hoc** |
| **6C** | **Day 1** | | 0.9096 | ns | 0.2707 | ns | 0.3085 | ns | N/A | Chi-square | |
| | **Day 2** | | 0.4623 | ns | 0.4623 | ns | >0.9999 | ns | | | |
| | **Day 3** | | 0.1915 | ns | 0.0982 | ns | 0.7047 | ns | | | |
| | **Day 4** | | 0.0479 | $ | 0.0985 | ns | 0.6988 | ns | | | |
| | **Day 5** | | 0.0479 | $ | 0.6358 | ns | 0.0943 | ns | | | |
| | | | **CTRL+Veh vs. SE+Veh ($)** | | **CTRL+Veh vs. SE+GP1a (#)** | | **SE+Veh vs. SE+GP1a (*)** | | **Normality ?** | **Test** | **Post-hoc** |

| | | | | | | | | | | |
|---|---|---|---|---|---|---|---|---|---|---|
| 6D | **Day 1** | 0.0640 | ns | 0.0617 | ns | 0.9947 | ns | Yes | Two-way repeated measures ANOVA | Tukey's |
| | **Day 2** | 0.0252 | $ | 0.0909 | ns | 0.7681 | ns | | | |
| | **Day 3** | 0.0283 | $ | 0.1591 | ns | 0.3617 | ns | | | |
| | **Day 4** | 0.01166 | $ | 0.0223 | # | 0.5866 | ns | | | |
| | | **CTRL+Veh** | | **SE+Veh** | | **SE+GP1a** | | | | |
| | **D1 vs. D3** | 0.3063 | ns | 0.2874 | ns | 0.0026 | ** | | | |
| | **D1 vs. D4** | 0.5753 | ns | 0.3625 | ns | 0.0158 | * | | | |
| | | **CTRL+Veh vs. SE+Veh ($)** | | **CTRL+Veh vs. SE+GP1a (#)** | | **SE+Veh vs. SE+GP1a (*)** | | **Normality ?** | **Test** | **Post-hoc** |
| 6E | **Day 1** | 0.0749 | ns | 0.1492 | ns | 0.7047 | ns | N/A | Chi-square | |
| | **Day 2** | 0.0479 | $ | 0.3562 | ns | 0.2248 | ns | | | |
| | **Day 3** | 0.0213 | $ | 0.3451 | ns | 0.0654 | ns | | | |
| | **Day 4** | 0.0441 | $ | >0.9999 | ns | 0.0308 | * | | | |
| | | **CTRL+Veh vs. SE+Veh** | | **CTRL+Veh vs. SE+GP1a** | | **SE+Veh vs. SE+GP1a** | | **Normality ?** | **Test** | **Post-hoc** |
| 6F | **Probe 1** | 0.0188 | * | 0.6546 | ns | 0.3534 | ns | No | Kruskall Wallis | Dunn's |
| 6G | **Probe 1** | 0.0045 | ** | 0.4507 | ns | 0.2306 | ns | No | Kruskall Wallis | Dunn's |
| 6H | **Probe 2** | 0.0464 | * | 0.9981 | ns | 0.0357 | * | No | Kruskall Wallis | Dunn's |
| 6I | **Probe 2** | 0.1972 | ns | 0.7983 | ns | 0.4817 | ns | No | Kruskall Wallis | Dunn's |
| | | **CTRL+Veh** | | **SE+Veh** | | **SE+GP1a** | | **Norm. ?** | **Test** | **Post-hoc** |
| 6K | **A vs. A'** | 0.3488 | ns | 0.6205 | ns | 0.9508 | ns | Yes | Two-way RM ANOVA | Šídák's |
| 6M | **A vs. B** | 0.0110 | * | 0.9971 | ns | 0.0022 | ** | | | |
| | | **CTRL+Veh vs. SE+Veh** | | **CTRL+Veh vs. SE+GP1a** | | **SE+Veh vs. SE+GP1a** | | **Normality ?** | **Test** | **Post-hoc** |
| 6N<br>6O | **"Group" factor** | 0.0039 | ** | 0.6280 | ns | 0.0385 | * | Yes | Two-way RM ANOVA | |
| 6P | **10 min** | 0.0080 | ** | 0.7010 | ns | 0.0452 | * | Yes | One-way ANOVA | Tukey's |

#### Figure 7

| | | **p-value** | | | | | | **Normality ?** | **Test** | **Post-hoc** |
|---|---|---|---|---|---|---|---|---|---|---|
| | | **CTRL+Veh vs. SE+Veh** | | **CTRL+Veh vs. SE+GP1a** | | **SE+Veh vs. SE+GP1a** | | | | |
| 7B | **O-maze** | 0.0257 | * | >0.9999 | ns | 0.0151 | * | No | Kruskall Wallis | Dunn's |
| 7D | **WET** | 0.7191 | ns | 0.3975 | ns | 0.0187 | * | No | Kruskall Wallis | Dunn's |

#### Figure 8

| | | **p-value** | | **Normality ?** | **Test** |
|---|---|---|---|---|---|
| 8A | **SE+Veh vs. SE+GP1a** | 0.0116 | * | N/A | Gehan-Breslow-Wilcoxon |
| 8B | **SE+Veh vs. SE+GP1a** | 0.0087 | ** | No | Mann-Whitney |

#### Figure S2

| | | | | | | |
|---|---|---|---|---|---|---|
| S2C | **CTRL+Veh vs. SE+Veh** | <0.0001 | **** | Yes | One-way ANOVA | Tukey's |
| | **CTRL+Veh vs. SE+GP1a** | 0.0916 | ns | | | |
| | **SE+Veh vs. SE+GP1a** | 0.0029 | ** | | | |

#### Figure S4

| | | | **p-value** | | **Normality ?** | **Test** | **Post-hoc** |
|---|---|---|---|---|---|---|---|
| S4A | Day 0 | **CTRL+Veh vs. SE+Veh** | >0.9999 | ns | No | Kruskall Wallis | Dunn's |
| | | **CTRL+Veh vs. SE+GP1a** | >0.9999 | ns | | | |
| | | **SE+Veh vs. SE+GP1a** | >0.9999 | ns | | | |
| S4B | Day 6 | **CTRL+Veh vs. SE+Veh** | <0.0001 | **** | Yes | One-way ANOVA | Tukey's |
| | | **CTRL+Veh vs. SE+GP1a** | 0.0161 | * | | | |
| | | **SE+Veh vs. SE+GP1a** | 0.0055 | ** | | | |
| S4C | Day 1 | **CTRL+Veh vs. SE+Veh ($)** | <0.0001 | $$$$ | Yes | Two-way ANOVA | Tukey's |
| | | **CTRL+Veh vs. SE+GP1a (#)** | <0.0001 | #### | | | |
| | | **SE+Veh vs. SE+GP1a (*)** | 0.0433 | * | | | |
| | Day 2 | **CTRL+Veh vs. SE+Veh ($)** | <0.0001 | $$$$ | | | |
| | | **CTRL+Veh vs. SE+GP1a (#)** | <0.0001 | #### | | | |
| | | **SE+Veh vs. SE+GP1a (*)** | 0.7445 | ns | | | |
| | Day 3 | **CTRL+Veh vs. SE+Veh ($)** | <0.0001 | $$$$ | | | |
| | | **CTRL+Veh vs. SE+GP1a (#)** | <0.0001 | #### | | | |
| | | **SE+Veh vs. SE+GP1a (*)** | 0.2587 | ns | | | |
| | Day 6 | **CTRL+Veh vs. SE+Veh ($)** | <0.0001 | $$$$ | | | |
| | | **CTRL+Veh vs. SE+GP1a (#)** | 0.0003 | ### | | | |
| | | **SE+Veh vs. SE+GP1a (*)** | 0.0315 | * | | | |

#### Figure S5

| | **p-value** | | **Test** | **Post-hoc** |
|---|---|---|---|---|

| | | | CTRL+Veh vs. SE+Veh ($) | | CTRL+Veh vs. SE+GP1a ($) | | SE+Veh vs. SE+GP1a (*) | | Normality ? | | |
|---|---|---|---|---|---|---|---|---|---|---|---|
| S5A | GFAP mRNA | SE+1D | <0.0001 | $$$$ | <0.0001 | $$$$ | 0.6565 | ns | Yes | One-way ANOVA | Tukey's |
| | | SE+9D | 0.0449 | $ | 0.0283 | $ | >0.9999 | ns | No | Kruskall Wallis | Dunn's |
| S5B | GFAP+ Area | SE+1D | 0.4737 | ns | 0.6359 | ns | 0.9576 | ns | Yes | Two-way ANOVA | Tukey's |
| | | SE+9D | 0.5273 | ns | 0.2715 | ns | 0.8995 | ns | | | |
| S5E | Iba1 mRNA | SE+1D | 0.2458 | ns | 0.3060 | ns | 0.9882 | ns | Yes | Two-way ANOVA | Tukey's |
| | | SE+9D | <0.0001 | $$$$ | <0.0001 | $$$$ | 0.9882 | ns | | | |
| S5F | Iba1+ Area | SE+1D | 0.2458 | ns | 0.3060 | ns | 0.9882 | ns | Yes | Two-way ANOVA | Tukey's |

#### Figure S6

| | | | | p-value | | Normality ? | Test | Post-hoc |
|---|---|---|---|---|---|---|---|---|
| S6D | IL-1β | Microglia | CTRL+Veh vs. SE+Veh | >0.9999 | ns | No | Kruskall Wallis | Dunn's |
| | | | CTRL+Veh vs. SE+GP1a | >0.9999 | ns | | | |
| | | | SE+Veh vs. SE+GP1a | >0.9999 | ns | | | |
| | | Mo-mΦs | SE+Veh vs. SE+GP1a | >0.9999 | ns | No | Mann-Whitney | |
| | | Other | CTRL+Veh vs. SE+Veh | 0.8756 | ns | No | Kruskall Wallis | Dunn's |
| | | | CTRL+Veh vs. SE+GP1a | 0.9303 | ns | | | |
| | | | SE+Veh vs. SE+GP1a | 0.9904 | ns | | | |
| | | CTRL+Veh | Microglia vs. Other | <0.0001 | **** | No | Mann-Whitney | |
| | | SE+Veh | Microglia vs. Mo-mΦs | 0.2261 | ns | Yes | One-way ANOVA | Tukey's |
| | | | Microglia vs. Other | 0.8507 | ns | | | |
| | | | Mo-mΦs vs. Other | 0.1126 | ns | | | |
| | | SE+GP1a | Microglia vs. Mo-mΦs | 0.5391 | ns | No | Kruskall Wallis | Dunn's |
| | | | Microglia vs. Other | >0.9999 | ns | | | |
| | | | Mo-mΦs vs. Other | 0.0760 | ns | | | |
| S6E | IL-6 | Microglia | CTRL+Veh vs. SE+Veh | 0.3032 | ns | No | Kruskall Wallis | Dunn's |
| | | | CTRL+Veh vs. SE+GP1a | 0.5391 | ns | | | |
| | | | SE+Veh vs. SE+GP1a | >0.9999 | ns | | | |
| | | Mo-mΦs | SE+Veh vs. SE+GP1a | 0.9715 | ns | Yes | Unpaired t-test | |
| | | Other | CTRL+Veh vs. SE+Veh | 0.2594 | ns | No | Kruskall Wallis | Dunn's |
| | | | CTRL+Veh vs. SE+GP1a | 0.1872 | ns | | | |
| | | | SE+Veh vs. SE+GP1a | >0.9999 | ns | | | |
| | | CTRL+Veh | Microglia vs. Other | >0.9999 | ns | No | Mann-Whitney | |
| | | SE+Veh | Microglia vs. Mo-mΦs | >0.9999 | ns | No | Kruskall Wallis | Dunn's |
| | | | Microglia vs. Other | 0.6991 | ns | | | |
| | | | Mo-mΦs vs. Other | 0.1579 | ns | | | |
| | | SE+GP1a | Microglia vs. Mo-mΦs | >0.9999 | ns | Yes | One-way ANOVA | Tukey's |
| | | | Microglia vs. Other | 0.0044 | ** | | | |
| | | | Mo-mΦs vs. Other | 0.0044 | ** | | | |
| S6F | TNFα | Microglia | CTRL+Veh vs. SE+Veh | 0.0626 | ns | No | Kruskall Wallis | Dunn's |
| | | | CTRL+Veh vs. SE+GP1a | 0.6154 | ns | | | |
| | | | SE+Veh vs. SE+GP1a | 0.7300 | ns | | | |
| | | Mo-mΦs | SE+Veh vs. SE+GP1a | 0.6073 | ns | Yes | Unpaired t-test | |
| | | Other | CTRL+Veh vs. SE+Veh | 0.8902 | ns | No | Kruskall Wallis | Dunn's |
| | | | CTRL+Veh vs. SE+GP1a | 0.1579 | ns | | | |
| | | | SE+Veh vs. SE+GP1a | 0.9519 | ns | | | |
| | | CTRL+Veh | Microglia vs. Other | <0.0001 | **** | No | Mann-Whitney | |
| | | SE+Veh | Microglia vs. Mo-mΦs | 0.0028 | ** | Yes | One-way ANOVA | Tukey's |
| | | | Microglia vs. Other | 0.0011 | ** | | | |
| | | | Mo-mΦs vs. Other | 0.5603 | ns | | | |
| | | SE+GP1a | Microglia vs. Mo-mΦs | <0.0001 | **** | Yes | One-way ANOVA | Tukey's |
| | | | Microglia vs. Other | <0.0001 | **** | | | |
| | | | Mo-mΦs vs. Other | 0.0340 | * | | | |
| S6G | IL-10 | Microglia | CTRL+Veh vs. SE+Veh | >0.9999 | ns | No | Kruskall Wallis | Dunn's |
| | | | CTRL+Veh vs. SE+GP1a | >0.9999 | ns | | | |
| | | | SE+Veh vs. SE+GP1a | 0.4008 | ns | | | |
| | | Mo-mΦs | SE+Veh vs. SE+GP1a | 0.2948 | ns | Yes | Unpaired t-test | |
| | | Other | CTRL+Veh vs. SE+Veh | >0.9999 | ns | No | Kruskall Wallis | Dunn's |
| | | | CTRL+Veh vs. SE+GP1a | >0.9999 | ns | | | |
| | | | SE+Veh vs. SE+GP1a | 0.9519 | ns | | | |
| | | CTRL+Veh | Microglia vs. Other | 0.3333 | ns | No | Mann-Whitney | |
| | | SE+Veh | Microglia vs. Mo-mΦs | 0.0178 | * | Yes | One-way ANOVA | Tukey's |
| | | | Microglia vs. Other | 0.9916 | ns | | | |
| | | | Mo-mΦs vs. Other | 0.0155 | * | | | |
| | | SE+GP1a | Microglia vs. Mo-mΦs | 0.0354 | * | Yes | One-way ANOVA | Tukey's |

| | | | | | | | | |
|---|---|---|---|---|---|---|---|---|
| | | | **Microglia vs. Other** | 0.9389 | ns | | | |
| | | | **Mo-mΦs vs. Other** | 0.0237 | * | | | |
| **S6H** | **IL-13** | **Microglia** | **CTRL+Veh vs. SE+Veh** | 0.4081 | ns | No | Kruskall Wallis | Dunn's |
| | | | **CTRL+Veh vs. SE+GP1a** | 0.4081 | ns | | | |
| | | | **SE+Veh vs. SE+GP1a** | >0.9999 | ns | | | |
| | | **Mo-mΦs** | **SE+Veh vs. SE+GP1a** | 0.6411 | ns | Yes | Unpaired t-test | |
| | | **Other** | **CTRL+Veh vs. SE+Veh** | 0.4081 | ns | No | Kruskall Wallis | Dunn's |
| | | | **CTRL+Veh vs. SE+GP1a** | >0.9999 | ns | | | |
| | | | **SE+Veh vs. SE+GP1a** | 0.5473 | ns | | | |
| | | **CTRL+Veh** | **Microglia vs. Other** | 0.3333 | ns | No | Mann-Whitney | |
| | | **SE+Veh** | **Microglia vs. Mo-mΦs** | 0.5030 | ns | Yes | One-way ANOVA | Tukey's |
| | | | **Microglia vs. Other** | 0.8691 | ns | | | |
| | | | **Mo-mΦs vs. Other** | 0.2815 | ns | | | |
| | | **SE+GP1a** | **Microglia vs. Mo-mΦs** | 0.8756 | ns | Yes | One-way ANOVA | Tukey's |
| | | | **Microglia vs. Other** | 0.9303 | ns | | | |
| | | | **Mo-mΦs vs. Other** | 0.9904 | ns | | | |
| **S6I** | **Arg-1** | **Microglia** | **CTRL+Veh vs. SE+Veh** | 0.4702 | ns | No | Kruskall Wallis | Dunn's |
| | | | **CTRL+Veh vs. SE+GP1a** | 0.0920 | ns | | | |
| | | | **SE+Veh vs. SE+GP1a** | >0.9999 | ns | | | |
| | | **Mo-mΦs** | **SE+Veh vs. SE+GP1a** | 0.8143 | ns | Yes | Unpaired t-test | |
| | | **Other** | **CTRL+Veh vs. SE+Veh** | >0.9999 | ns | No | Kruskall Wallis | Dunn's |
| | | | **CTRL+Veh vs. SE+GP1a** | >0.9999 | ns | | | |
| | | | **SE+Veh vs. SE+GP1a** | >0.9999 | ns | | | |
| | | **CTRL+Veh** | **Microglia vs. Other** | 0.3333 | ns | No | Mann-Whitney | |
| | | **SE+Veh** | **Microglia vs. Mo-mΦs** | 0.0709 | ns | Yes | One-way ANOVA | Tukey's |
| | | | **Microglia vs. Other** | 0.9998 | ns | | | |
| | | | **Mo-mΦs vs. Other** | 0.0691 | ns | | | |
| | | **SE+GP1a** | **Microglia vs. Mo-mΦs** | 0.0011 | ** | Yes | One-way ANOVA | Tukey's |
| | | | **Microglia vs. Other** | 0.9942 | ns | | | |
| | | | **Mo-mΦs vs. Other** | 0.0010 | *** | | | |
